# The inflammasome adaptor protein ASC present differential intracellular and extracellular functions to trigger immunometabolic dysregulation and dysbiosis during obesity

**DOI:** 10.1101/2025.01.16.633385

**Authors:** Juan José Martinez-Garcia, Sara Rodríguez, Adriana Guijarro, Ignacio Quevedo-Romero, Sandra V. Mateo, Cristina Molina-López, Laura Hurtado-Navarro, Alberto Baroja-Mazo, Pablo Pelegrin

## Abstract

The inflammasomes are a component of the innate immune system that induce inflammation after activation by danger and pathogen signals, this process is of extreme importance in the gut, where high-saturated fat diets modulate inflammasome activity resulting in dysbiosis and inflammation. In this work, we found that the common inflammasome adaptor protein, apoptosis speck-like protein with a caspase activation domain (ASC), is essential for the development of metabolic dysregulation during high fat diet. ASC-deficient mice (*Pycard*^-/-^) presented lower weight gain and improved glucose tolerance, enhancing metabolism, and reducing inflammation in both intestine and liver. The extracellular administration of ASC oligomers to *Pycard*^-/-^ mice partially rescued dysbiosis, intestinal metabolism and inflammation. Therefore, ASC presents intracellular functions activating the inflammasome, as well as extracellular function as an oligomer during metabolic dysregulation, emerging ASC as a potential drug target beneficial to treat obesity and metabolism-associated liver diseases.

## Introduction

Nucleotide-binding domain leucine-rich repeat receptors (NLRs) constitute an important part of the inflammatory response, responding to a wide variety of danger and pathogens associated molecular patterns (1,2). NLRs, including the most studied NLRP3, are highly expressed in immune myeloid cells as well as in epithelial cells (3). NLRP3 and other NLRs can oligomerize forming inflammasomes that activate caspase-1 to mature the proinflammatory cytokines interleukin (IL)-1β and IL-18 (4,5). Caspase-1 also processes the protein gasdermin D (GSDMD), which N-terminal fragment form pores in the plasma membrane initiating the release of IL-1β and IL-18 and induces an immunogenic programmed cell death termed pyroptosis (6). During pyroptotic cell death the plasma membrane protein ninjurin-1 (NINJ1) induces membrane rupture (7). Most of the inflammasomes require interaction and oligomerization of the apoptosis-associated speck-like protein containing a caspase-1 activating domain (ASC) to activate caspase-1. Inflammasome oligomers of ASC are externalized to the extracellular milieu during pyroptosis to amplify the inflammatory response (8,9). Thus, beyond the intracellular scaffold function of ASC to recruit and activate caspase-1, extracellular ASC oligomers play a role in different pathologies amplifying the inflammatory response and causing amyloid deposition (10–13). Intestinal microbiota plays a crucial role contributing to the host’s health and survival (14). This makes the gut extremely sensitive to microbiota changes, which controls colonisation by opportunistic bacteria and triggers innate immune responses, including inflammation (15,16). Subsequently, the effect of the gut microbiome controlling the activity of the inflammasome has been reported with both antibiotics and germ-free (GF) conditions (17,18). The participation of the different inflammasomes, including the NLRP3, has been tightly associated with diet-related metabolic diseases, such as obesity and type II diabetes (19). In fact, the consume of diets with high saturated fats (HFD) induces an inflammatory response and increase IL-1β production (20). Furthermore, the NLRP6 inflammasome, which is present in epithelial intestinal cells, maintains gut microbiota homeostasis through the production of IL-18 (21). Furthermore, the participation of pyroptosis induced by GSDMD has been described as a protective agent that controls intestinal microbiota and repairs intestinal mucosa (22). However, there is controversy with the role of pyroptosis in obesity, as it is not only associated with an increase in body weight but is also capable of inducing metabolic dysregulations, such as blockage of cholesterol transport for subsequent catabolic processes, the accumulation of atheroma plaques, and the development of inflammatory diseases such as steatohepatitis (23–25). Since there is a clear involvement of different inflammasomes in metabolic dysregulation, the role of the common inflammasome adaptor protein ASC, and its intracellular and extracellular functions in this disease is not fully understood. In this study we focus on the effect of ASC deficiency on metabolic dysregulation induced by HFD and uncover a differential function of ASC, acting as intracellular scaffold to form inflammasome and its extracellular function as ASC oligomers. We identify that the deletion of ASC protects mice in response to a HFD, increase the tolerance to glucose and exacerbate glycolytic and aerobic metabolism in HFD conditions, with a decreased inflammatory response. ASC deficiency also affected gut epithelial integrity and permeability, regulating gut microbiome. Extracellular ASC oligomers administration in ASC-deficient mice partially revert dysbiosis, intestinal metabolism and inflammation, revealing a differential role of intracellular and extracellular ASC in metabolic dysregulation. This work demonstrates that ASC could be a potential therapeutic target to treat metabolic dysfunction associated to steatosis liver disease and type II diabetes.

## Results

### ASC deficiency prevents glucose intolerance under high-fat-diet conditions

We initially asked whether ASC deficiency can modulate weight gain in obesity after feeding with a HFD. Weight of *Pycard*^-/-^ was lower when compared to wild type from the third month of HFD, although the weight reduction was not statistically significant **(Figure 1A)**. To monitor potential weight changes during the time dependent on the inflammasome, weight from C57/BL6 background wild type (*Pycard*^+/+^) mice and *Pycard*^-/-^ mice from same age was monitored during 5 months in normal chow diet (NCD). We observed that the *Pycard*^-/-^ mice tended to gain less weight than the *Pycard*^+/+^ in both males and females, but this difference was not statistically significant **(Figure 1B)**. However, this tendency reveals that deficiency of ASC can prevent body weight gain, and this effect was independently on the sex **(Figure 1B)**. HFD-dependent weight gain was confirmed comparing HFD to NCD fed wild type mice **(Suppl. Figure 1A)**. We next analysed glucose tolerance by measuring the glucose concentration after intraperitoneal glucose administration in mice feed with HFD and observed an increase of glucose tolerance in the *Pycard*^-/-^ mice respect to wild type mice after 3 months of HFD **(Figure 1C)** and after 5 months of HFD **(Figure 1D)**. As expected, glucose intolerance was mediated by the hypercaloric diet, as HFD fed mice presented higher glucose intolerance compared to NCD fed mice **(Suppl. Figure 1B).** Additionally, we observed that *Pycard*^-/-^ mice fed with a normal diet tended to increase glucose tolerance in comparison with wild type mice **(Figure 1E)**. However, genotype-dependent differences in NCD fed mice were smaller than observed in HFD mice **(Figure 1C, D)**.

**Figure 1.**
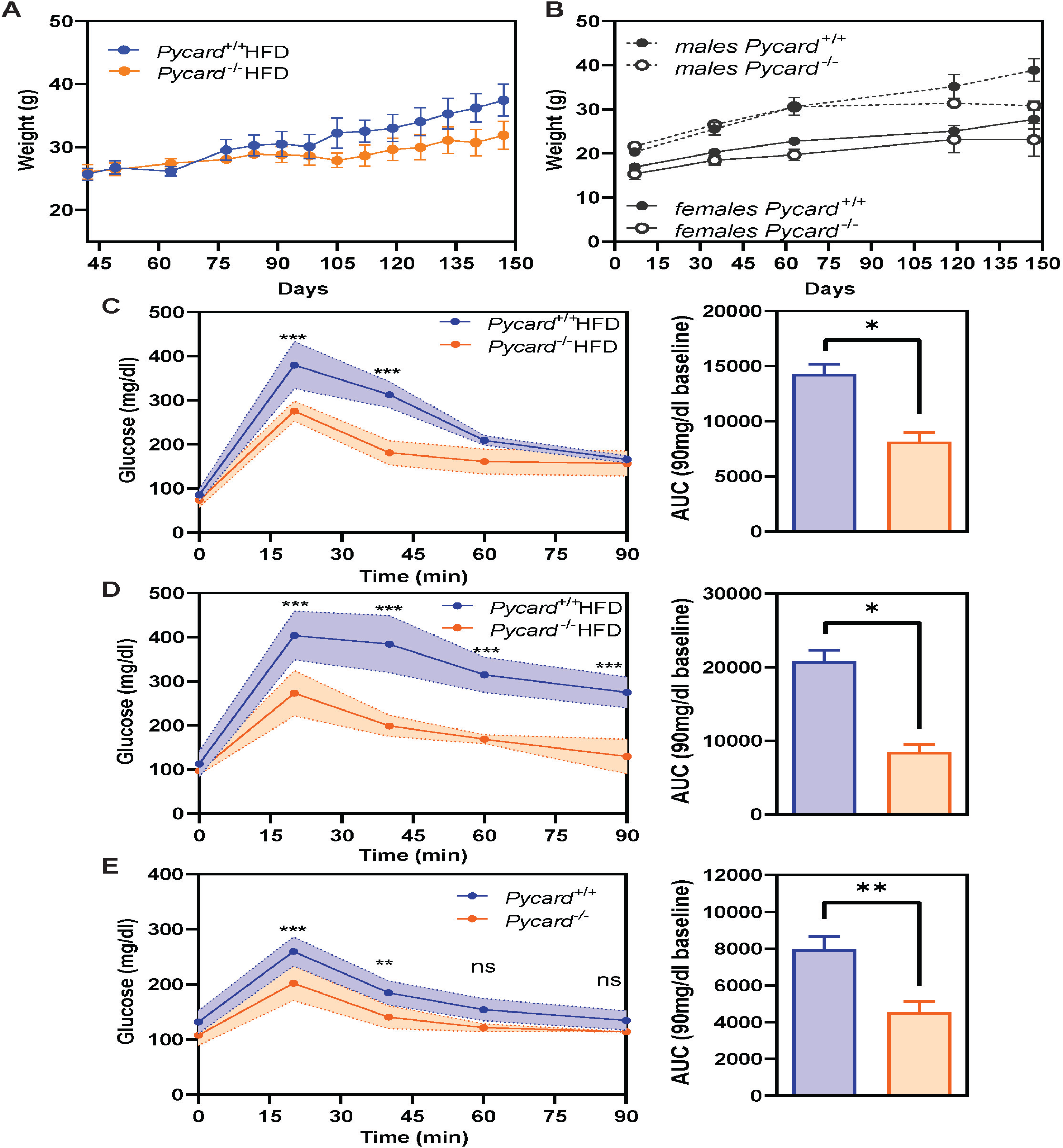
ASC deficiency prevents glucose intolerance during high fat diet. **A.** Body weight from wild type (*Pycard*^+/+^, blue, n=5), and *Pycard*^-/-^ (orange, n=4) mice fed with a high fat diet (HFD) for 5 months. **B.** Body weight from *Pycard*^+/+^ (balck, n=3-4), *Pycard*^-/-^ (white, n=2) mice separated by sex, males (discontinuous line) and females (continuous line) fed with a normal chow diet (NCD) for 5 months. **C-E.** Glucose tolerance (left) and area under the curve (AUC, right) from mice fed with HFD (C,D) or NCD (E) during 3 months (C,E) or 5 months (D). Wild type (*Pycard*^+/+^, blue, n=5), and *Pycard*^-/-^ (orange, n=4) mice. Bars or dots represent the mean, error bars represent SEM. Two-way ANOVA with multivariant Sidak’s test was used for all panels. Mann-Whitney’s test was done for C-E. (ns, p>0.05, *p≤0.05; **p≤0.01; *** p≤0.001).

### ASC controls peritoneal cell infiltration in high-fat-diet and mediates IL-1β release

We observed an increase of the total number of cells in the peritoneum upon 5 months of HFD respect NCD fed wild type mice **(Figure 2A)**. To investigate inflammasome function in the infiltrated peritoneal cells during HFD conditions, we cultured adherent peritoneal cells from mice and measure IL-1β in cell supernatants after *in vitro* NLRP3 inflammasome activation. Peritoneal cells from wild type mice fed with HFD presented NLRP3-dpenent IL-1β release just by LPS treatment **(Figure 2B)**, being potentiated by nigericin in both NCD and HFD conditions **(Figure 2B)**. This suggests increase sensitivity to activate IL-1β release in peritoneal infiltrated cells in mice fed with a HFD. After 5 months of HFD the *Pycard*^-/-^ mice presented a higher increase of peritoneal cells when compared to wild type **(Figure 2C)**, but as expected, IL-1β release in peritoneal cells from *Pycard*^-/-^ mice was decreased when compared to wild type cells **(Figure 2D)**. Absence of IL-1β release was due to the *Pycard*^-/-^ genotype and not by the HFD effect **(Suppl. Figure 1C)**. As control, we found that IL-6 release from peritoneal cells was dependent on LPS stimulation, being independent of the diet or the genotype **(Figure 2E,F, Suppl. Figure 1D)**. These data suggest that the absence of ASC could reduce inflammasome-dependent peritoneal cell death and IL-1β production and reduce obesity-induced inflammation.

**Figure 2.**
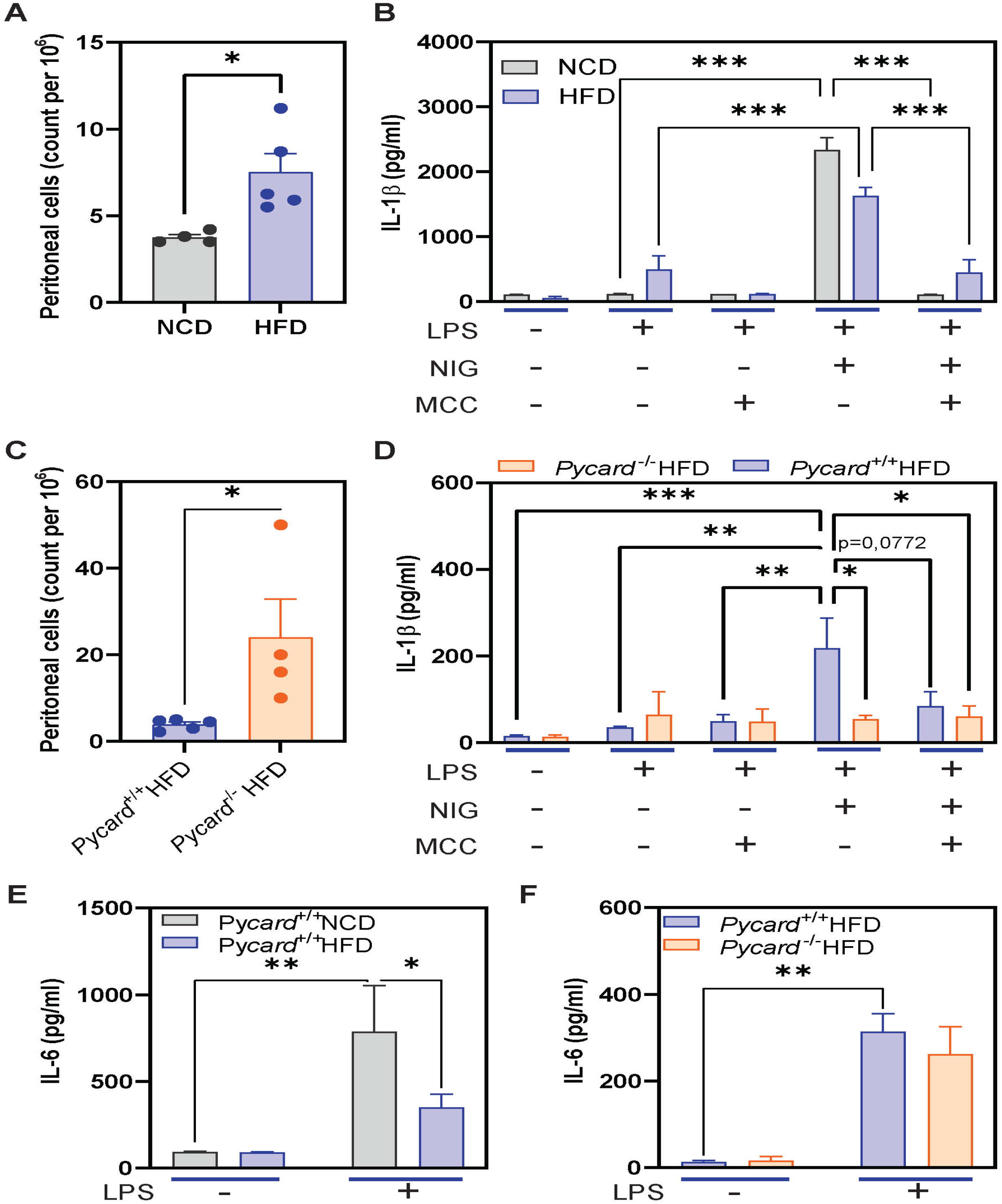
High fat diet increases intestinal permeability and reduces peritoneal cells dependent on ASC. **A, B**. Number of peritoneal cells (A) and interleukin (IL)-1β release from peritoneal cells (B) treated as indicated in the figure from wild type mice fed for 5 months with a normal chow diet (NCD, n=4, grey) or high fat diet (HFD, n=5, blue). **C, D**. Number of peritoneal cells (C) and IL-1β release from peritoneal cells (D) treated as indicated in the figure from wild type (*Pycard*^+/+^, n=5, blue) or *Pycard*^-/-^ (n=4, orange) mice fed for 5 months with a HFD. LPS: lipopolysaccharide; NIG: nigericin; MCC: MCC950. **E, F.** IL-6 release from peritoneal cells collected from mice treated as in A-D and then *in vitro* stimulated as described in the figure. Each dot in A,C represents a distinct animal, bars represent the mean, error bars represent SEM. Mann-Whitney’s test was used for A,C. Two-way ANOVA with multivariant Sidak’s test was used for B, D-F (ns, p>0,05; *p≤0.05; **p≤0.01; *** p≤0.001).

### ASC restricts gut microbiota biodiversity and controls intestinal permeability

We next wondered if ASC could influence gut dysbiosis induced by HFD. We determined 16S RNA in faeces and found that the microbiota composition changed in *Pycard*^-/-^ mice fed with HFD during 5 months in comparison with wild type mice **(Figure 3A-C)**. After HFD, wild type mice accumulated the 97% of the abundance on the four predominant class of bacteria: Clostridia, Bacilli, Verrucomicrobiae and Bacteroidia **(Figure 3A)**. Within the most abundant class of bacteria that was influenced by the deficiency of ASC under HFD, we encounter an increase of Clostridia, meanwhile Verrucomicrobiae class decreased on *Pycard*^-/-^ HFD mice **(Figure 3B).** Moreover, *Pycard*^-/-^ mice presented higher percentages of relative abundance for other less abundant class of bacteria, specifically on Coribacteria, Desulfovibrionia, and Deferribacteres **(Figure 3B)**. As a control, we observed in a different batch of mice that wild type with NCD presented low proportions of Verrucomicrobiota **(Suppl. Figure 2A)**, which strongly increased after HFD **(Suppl. Figure 2B)**. The increase diversity of bacteria classes found in the *Pycard*^-/-^ after HFD was also maintained where we analysed the different represented families of bacteria from the total microbiome **(Figure 3C)**. It is particularly striking the increase of Lachnospiraceae family in the *Pycard^-/-^* mice after HFD, meanwhile the mucin-degrading bacteria from the Akkermansiaceae family decreased **(Figure 3C)**. We corroborated in a second batch of mice that Lachnospiraceae family was increased in absence of ASC **(Suppl. Figure 3A)**. After cohousing between wild type and *Pycard*^-/-^ mice, genetic differences were abolished before and after the HFD treatment except for an increase of Lactobacillaceae in *Pycard^-/-^* mice after HFD **(Suppl. Figure 3B,C)**. Moreover, the average of the alpha-biodiversity Shannon Index increased in *Pycard^-/-^* mice after HFD **(Figure 3D)**. Within the most representative genus of bacteria, *Akkermansia* abundance tended to decrease in absence of ASC **(Figure 3E)**. In a different batch of mice, we tested the abundance of *Akkermansia* after a HFD in comparison with a NCD, where we observed that HFD highly increased the proportion of this bacterium **(Suppl. Figure 2C)**. Contrarily to *Akkermansia*, and accordingly with observed for class and families, other genus of bacteria strongly increased its abundance, becoming the most abundant bacteria in the microbiome of *Pycard^-/-^* mice after HFD, including *Odoribacter*, *Lachnospiraceae* NK4A136, *Coriobacteriaceae* UGC002, and *Lactobacillus* **(Figure 3F-I)**. On the contrary, ASC deficiency decreased the proportion of potential pathobionts such as *Streptococcus* and *Dorea* **(Figure 3J, K)**. We next evaluated intestinal permeability by FITC-dextran measurement on the plasma of wild type and *Pycard*^-/-^ mice fed with a NCD or a HFD and found that intestinal permeability increased in wild type mice after HFD, but not in *Pycard^-/-^* mice **(Figure 3L)**. We validated in a second batch of mice that HFD increased intestinal permeability in comparison with NCD fed mice **(Suppl. Figure 2D)**. These data show that ASC deficiency prevent dysbiosis in response to HFD, maintaining a low intestinal permeability, probably due to the decrease of pathobionts and mucin-degrading bacteria that influence intestinal epithelial protective functions.

**Figure 3.**
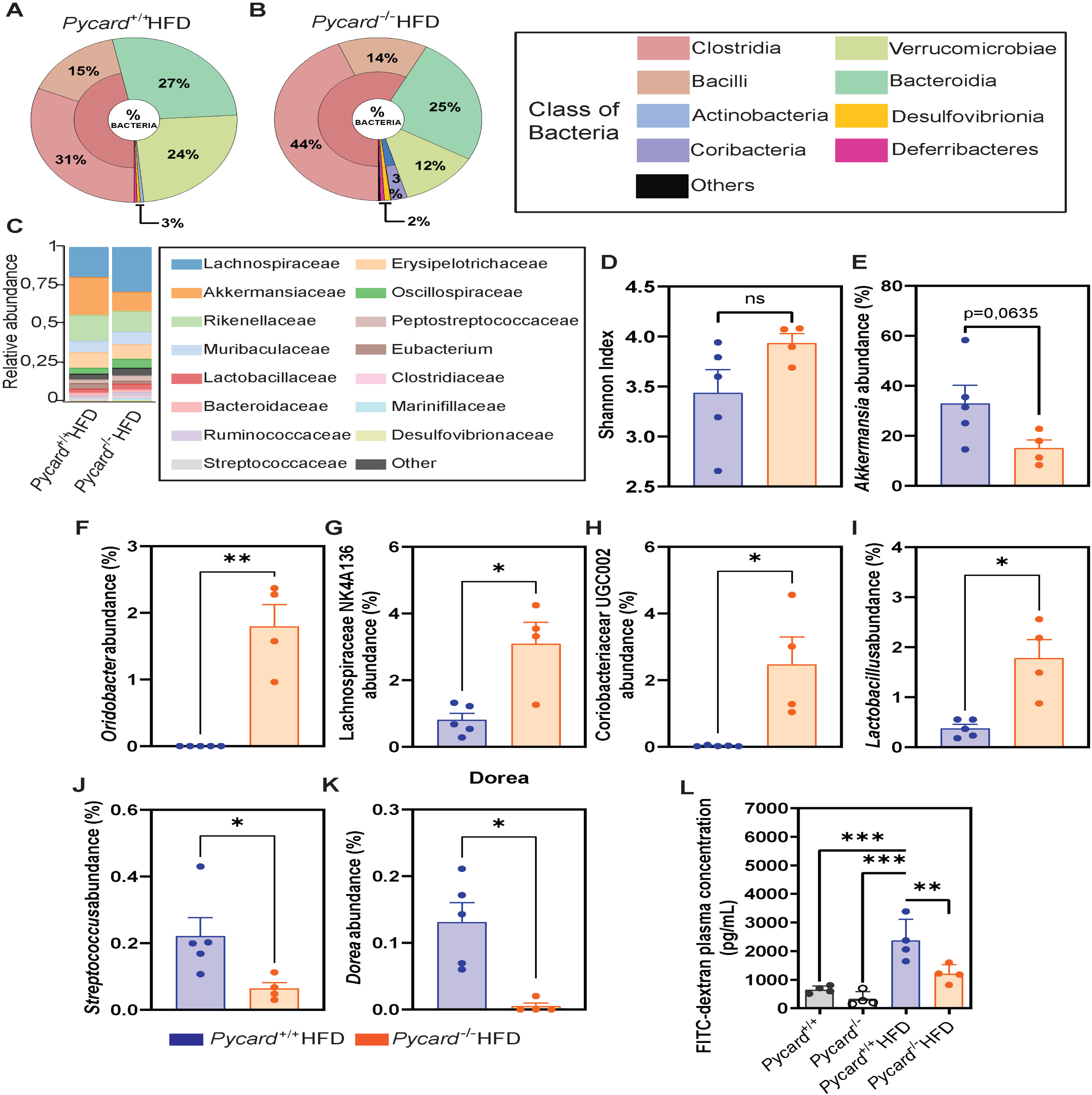
ASC modulate microbiota biodiversity during high fat diet. **A-C.** Relative microbiota abundance at level of class (A, B) or families (C) on wild type (*Pycard*^+/+^, n=5) (A, C) or *Pycard*^-/-^ mice (n=4) (B, C) after 5 months of high fat diet (HFD). **D**. Shannon index from the same conditions as in A-C. **E-K**. Relative abundance of *Akkermansia* (E), *Odoribacter* (F), Lachnospiraceae NK4A136 (G), Coriobacteriaceae UGC002 (H), *Lactobacillus* (I), *Streptococcus* (J), and *Dorea* (K) in the same conditions as in A-C. **L.** Intestinal permeability measured by 4 kDa FITC-dextran in *Pycard*^+/+^ NCD (grey, n=4), *Pycard*^+/+^ HFD (blue, n=5), *Pycard*^-/-^ NCD (white, n=4), and *Pycard*^-/-^ HFD (orange, n=4) after 5 months of high fat diet (HFD). Each dot represents a distinct animal, bars represent the mean, error bars represent SEM. Mann-Whitney’s test was done for D-K. One-way ANOVA with multivariant Holm-Sidak’s test was used for L (ns p>0,05; *p≤0.05; **p≤0.01; *** p≤0.001).

### ASC facilitates bacteria attachment to intestinal villi during high-fat-diet

Since the absence of ASC reduces intestinal permeability and prevents dysbiosis, we next examined if the intestinal mucosa was altered after HFD. We observed that wild type mice fed with HFD presented severe bacterial colonization in the small intestinal *villi* **(Figure 4A)**. Additionally, exacerbated amount of mucin was observed in the lumen of the intestinal tissue, meanwhile the space between *villi* presented numerous areas without mucin **(Figure 4A).** Contrarily, bacteria attached to the *villi* decreased in *Pycard*^-/-^ mice, and mucin amount was controlled in the border of each intestinal *villi* and uniformly disposed **(Figure 4B)**. As a control, when the intestinal *villi* of wild type mice fed with NCD were compared with the pathologic phenotype of the HFD fed mice, we observed a normal intestinal histology with low bacteria in the *villi* **(Suppl. Figure 2E, F)**. Regarding the uncontrolled amount of mucin found in the small intestine of wild type mice, we wondered if the absence of ASC would trigger a transcriptional profile to protect mucin degradation from bacteria. Bulk RNA sequencing in the small intestine of mice fed with HFD showed increasing transcript levels of the β-galactoside sialyltransferases encoding genes (*ST6Galnac2-6*) in *Pycard*^-/-^ mice **(Figure 4C)**, which could explain the protection of mucin between *villi*, being less accessible to mucin-degrading bacteria.

**Figure 4.**
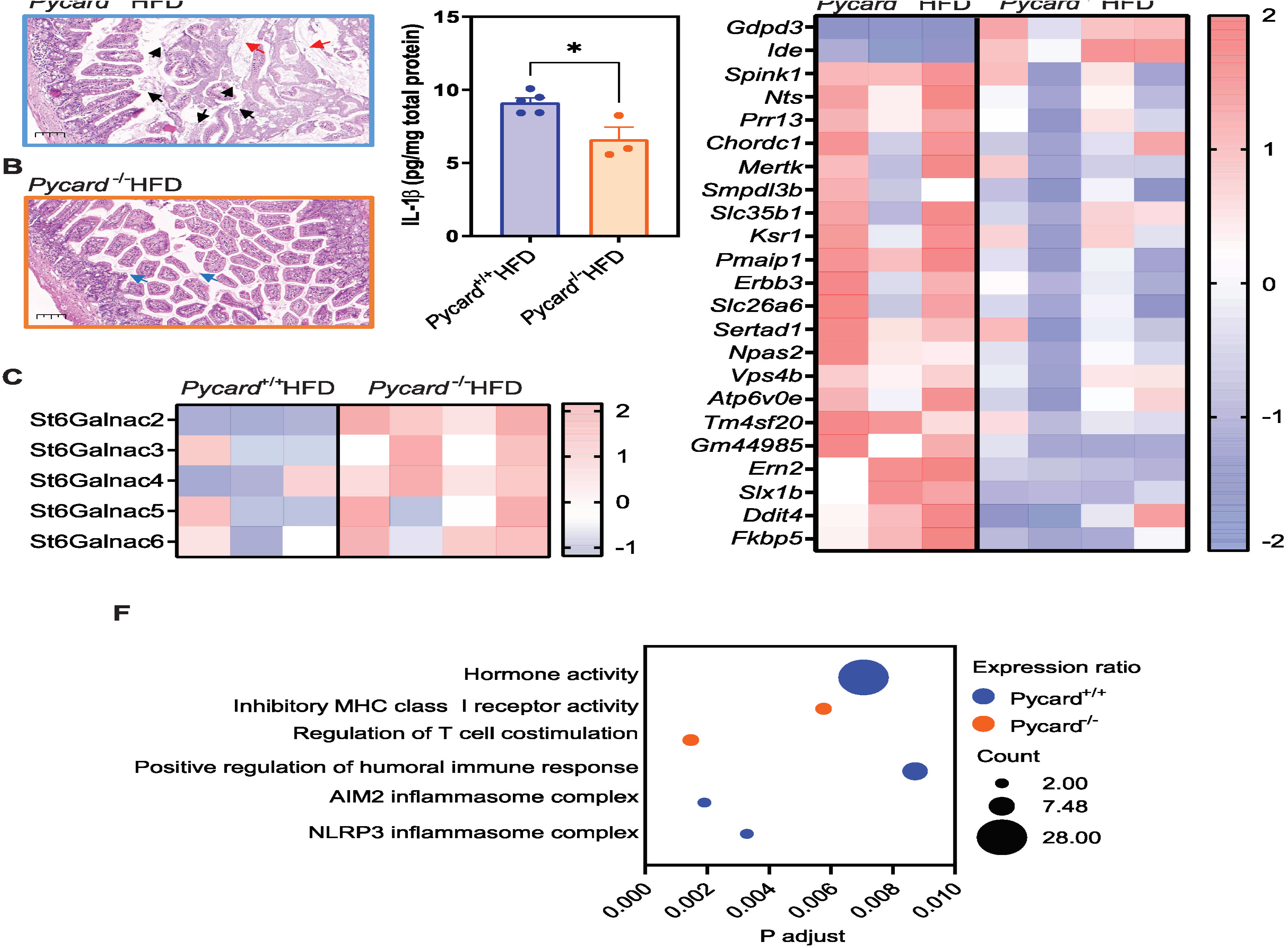
ASC induces bacteria attachment in small intestinal *villi* during high fat diet. **A, B.** Haematoxylin and eosin histology of the small intestine of wild type *Pycard*^+/+^ (A) and *Pycard*^-/-^ (B) mice fed for 5 months with a high fat diet (HFD). Scale bar: 200 μm; red arrows denote not attached mucus in the intestinal lumen; black arrows indicate presence of bacteria; blue arrows indicate presence of mucus between *villi*. **C.** *ST6Galnac* gene family transcript expression in the small intestine of mice treated as in A, B. **D.** Interleukin (IL)-1β in the small intestine of mice treated as in A, B. Wild type (*Pycard*^+/+^, n=5, blue), and *Pycard*^-/-^ (n=3, orange) mice. **E.** Differential expressed genes in the small intestine of mice treated as in A, B. **F.** Gene Ontology differential expressed pathways in the small intestine of mice treated as in A, B. Wild type (*Pycard*^+/+^, n=3), and *Pycard*^-/-^ (n=4,) mice. Each dot represents a distinct animal, bars represent the mean, error bars represent SEM. Mann-Whitney’s test was used for D (*p≤0.05).

Regarding the importance of the ASC-dependent inflammasomes in glucose tolerance **(Figure 1C,D)**, we measured the amount of IL-1β in the small intestine, which was decreased in *Pycard*^-/-^ feed with a HFD compared to wild type mice **(Figure 4D)**. Bulk RNA sequencing demonstrated that the presence of ASC increased the expression of genes related to intestinal inflammation (e.g. *Mertk, Smpdl3b, Fkbp5)*, epithelial integrity and mucus secretion (e.g. *Spink1, Pmaip1, Erbb3, Sl26a6, Ern2)*, and epithelial proliferation and regeneration (e.g. *Ksr, Npas2, Sertad1, Slx1b, Ddit)* **(Figure 4E)**. However, the *Pycard*^-/-^ mice have increased expression of metabolic and anti-inflammatory transcript genes (*Gdpd3* and *Ide*) **(Figure 4E).** Gene Ontology (GO) analysis from the differential expressed genes in the small intestine revealed that meanwhile the most enriched pathways in the HFD fed *Pycard*^-/-^ mice were the regulation of the T cell response and the regulation of the MHC class I and II activity, on *Pycard*^+/+^ mice, the AIM2 and NLRP3 inflammasome, hormonal activity and positive regulation of the humoral immune response were between the top 10 highly expressed pathways **(Figure 4F, Suppl. Figure 4)**. All together the deficiency of ASC induced in the small intestine mucosa a protective mechanism during HFD, reducing inflammation, increasing mucosal protection and preventing bacteria attachment to the *villi*.

### ASC deficiency increases glycolytic and mitochondrial metabolism in the small intestine during obesity

As the RNA sequencing revealed a tendency to increase some metabolic genes under HFD treatment, which could also explain the elevation of IL-1β, we then analyzed in the small intestine the expression of key metabolic genes related to glycolysis and the tricarboxylic acids (TCA) cycle. Interestingly, we found that the glycolytic *Pkm* gene coding for the pyruvate kinase enzyme was highly expressed in the *Pycard*^-/-^ mice after HFD **(Figure 5A)**. Contrarily, the anaerobic respiration-related gene *Ldha*, corresponding to the enzyme lactate dehydrogenase, decreased in the *Pycard*^-/-^ mice **(Figure 5B)**. Consequently, we observed an increase in the expression of *Pdha* gene, which encodes the pyruvate dehydrogenase enzyme **(Figure 5C)**. This observation could indicate that the high glycolytic state of the small intestine in the *Pycard*^-/-^ mice after HFD drives to an aerobic metabolism, and not to the anaerobic pathway. To verify this switch, we also analyzed the expression of enzymes implicated in mitochondrial metabolism in the small intestine, including those for the TCA cycle, the oxidation of fatty acids and the catabolism of amino acids. We found that the expression of *Nnt*, *Glud1* and *Sdha* genes increased in the *Pycard*^-/-^ mice after HFD **(Figure 5D-F)**. We also demonstrated that the expression of the β-oxidation-related gene *Cpt1a* increased in *Pycard*^-/-^ mice respect to wild type mice **(Figure 5G)**. These results align with the fact that the *Pycard*^-/-^ have decreased weight and increased glucose tolerance independently on the HFD probably due to a high catabolism and energy uptake.

**Figure 5.**
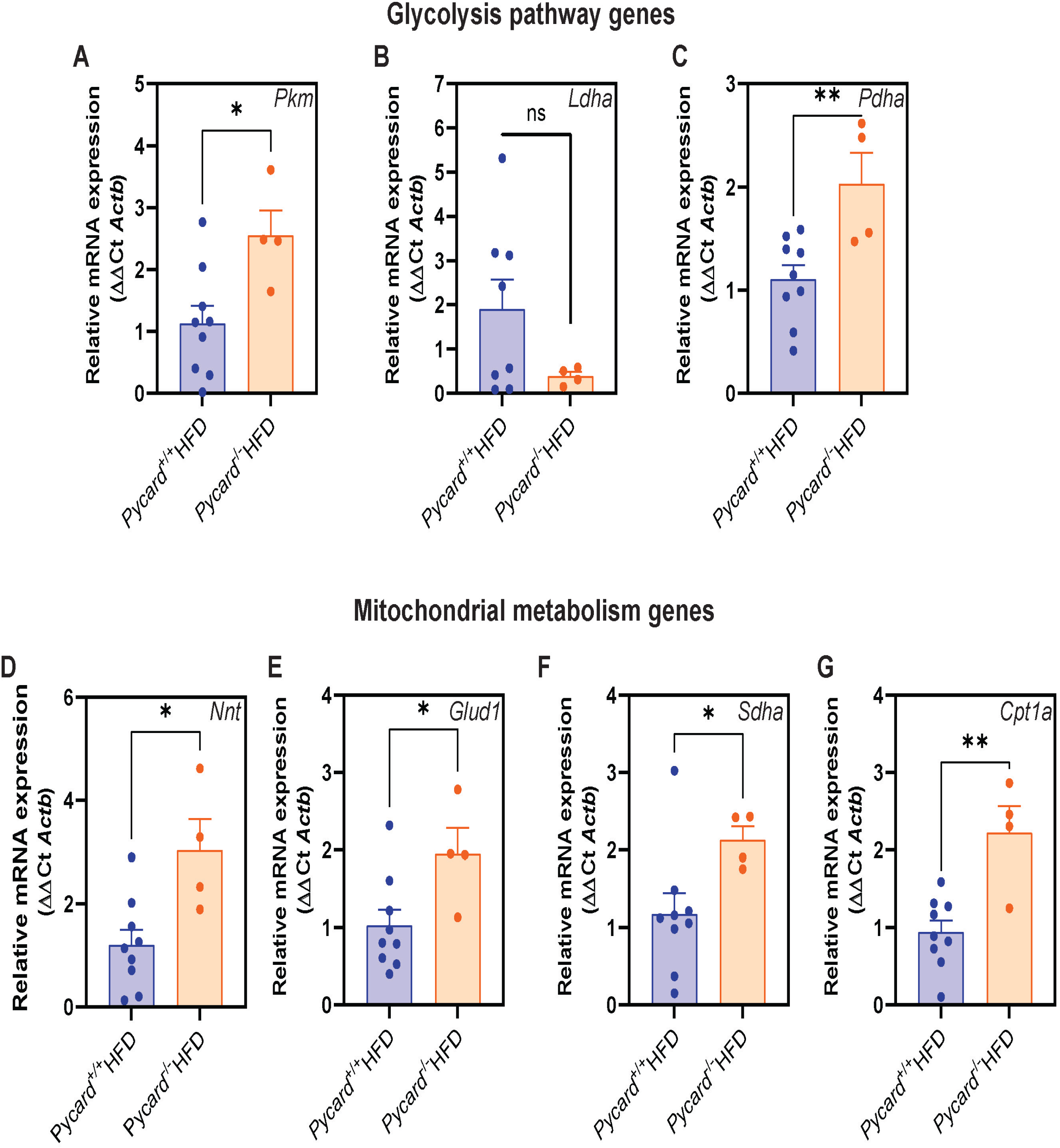
ASC deficiency increases glycolytic and mitochondrial metabolism related genes during high fat diet. **A-G.** Relative mRNA expression of *Pkm* (A), *Ldha* (B), *Pdha* (C), *Nnt* (D), *Glud1* (E), *Sdha* (F), and *Cpt1a* (G) on the small intestine of wild type (*Pycard*^+/+^, n=10, red) and *Pycard*^-/-^ (n=4, green) mice fed with HFD for 5 months. Bars represent the mean; error bars represent SEM. Unpaired *t*-test was used (ns, p>0.05; *p≤0.05; **p≤0.01).

### ASC deficiency enhances hepatic metabolism and reduces inflammation and fat accumulation under high-fat-diet

We then investigated if *Pycard*^-/-^ mice have an altered liver physiopathology and metabolism during HFD. Initially we observed that wild type mice fed with HFD presented an exacerbated fat accumulation in the liver, with some isolated inflammatory foci **(Figure 6A)**. Contrarily, the accumulation of fat in the liver was absent in ASC-deficient mice after HFD **(Figure 6B)**. The accumulation of hepatic fat due to the HFD in wild type was corroborated when compared with NCD fed mice **(Suppl. Figure 5A, B)**. Similarly to small intestinal mucosa, in the liver there was also decreased IL-1β production in the *Pycard*^-/-^ mice fed with HFD **(Figure 6C),** and no changes between genotypes were observed for IL-6 **(Figure 6D).** The high hepatic increase of the inflammasome-dependent cytokine IL-1β, was specific for the HFD, as NCD mice for both genotypes, showed basal levels of IL-1β **(Suppl. Figure 5C, D)**. Moreover, we observed increased expression of metabolic genes in the liver of *Pycard*^-/-^ HFD fed mice, in particular genes related with the PPARα signaling pathway, as the master energy homeostasis regulator *Pgc1a* **(Figure 6E)**. Similar to the small intestine, the mitochondrial metabolic genes *Nnt*, *Glud1* and *Sdha* were more expressed in the *Pycard*^-/-^ mice after HFD **(Figure 6F-H)**, and the fatty acid oxidation-related gene *Cpt1a* slightly tended to increase in the absence of ASC **(Figure 6I)**. Interestingly, we observed a marked increased expression of the ketone bodies-dependent gene *Hmgcs2* in *Pycard*^-/-^ mice fed with a HFD **(Figure 6J)**. Overall, ASC is important for the development of the metabolic syndrome induced by HFD.

**Figure 6.**
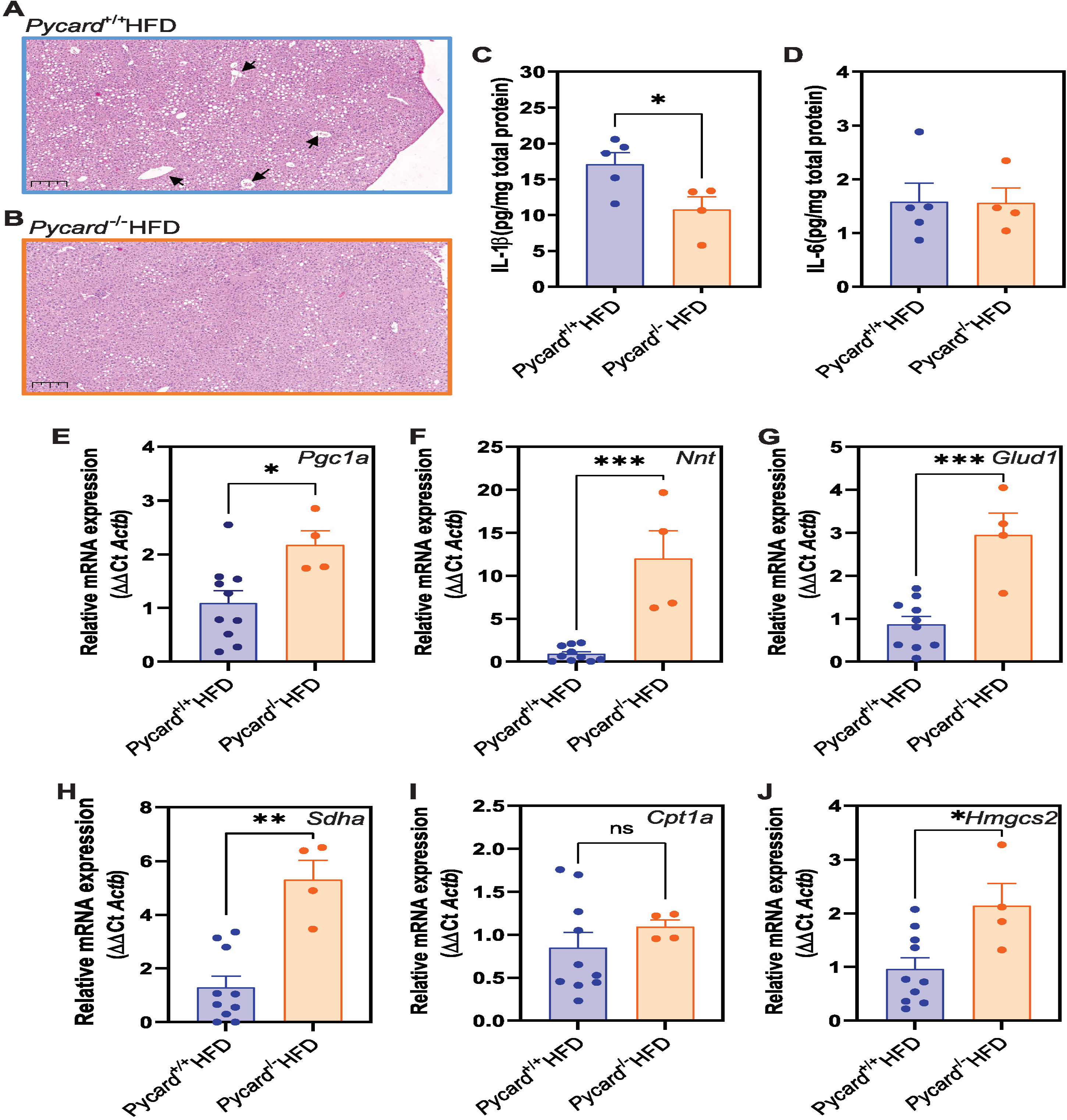
ASC decreases hepatic metabolism and enhances inflammation under high fat diet. **A, B.** Haematoxylin and eosin histology of the liver of wild type (*Pycard*^+/+^, A) and *Pycard*^-/-^ (B) mice fed for 5 months with a high fat diet (HFD). Scale bar: 200 μm; Arrows indicate hepatic vascularization. **C, D.** Interleukin (IL)-1β (C) and IL-6 (D) in the liver of mice treated as in A-B. **E-J.** Relative mRNA expression of *Pgc1a* (E)*, Nnt* (F)*, Glud1* (G)*, Sdha* (H)*, Cpt1a* (I) and *Hmgcs2* (J) in the liver of mice treated as in A-B. For C-J, *Pycard*^+/+^ mice (n=10) are represented in blue and *Pycard*^-/-^ mice (n=4) are represented in orange. Each dot in C,D represents a distinct animal, bars represent the mean, error bars represent SEM. Unpaired *t*-test was used for C-J (ns, p>0.05; *p≤0.05; *** p≤0.001).

### Extracellular ASC oligomers partially control metabolic syndrome during obesity

Beyond intracellular function of ASC forming inflammasomes, ASC oligomeric particles could also reach the extracellular milieu after pyroptosis being important for different pathologies (8–13,26). To explore if extracellular ASC oligomers were important for the development of metabolic syndrome after HFD, we rescued the *Pycard*^-/-^ mice with intraperitoneal administration of recombinant ASC oligomers **(Suppl. Figure 6A)**. Intestinal permeability was slightly reverted by administration of recombinant ASC oligomers, but differences were not significant comparing to the *Pycard*^+/+^ positive control **(Figure 7A)**. The administration of recombinant ASC oligomers produced a change in the gut microbiota composition of *Pycard*^-/-^, which experimented a rescue to an intermediate microbiome phenotype between *Pycard*^+/+^ and *Pycard*^-/-^ mice **(Figure 7B)**. These changes included the disappearance of some important beneficial bacteria genus found in *Pycard*^-/-^ mice fed with a HFD, such as *Lactobacillus* and *Coriobacteriaceae* UGC002, as well as less abundant bacteria such as *Enteractinococcus* and *Atopostipes* **(Figure 7B)**. Meanwhile, other bacteria such as *Oridobacter* and *Lachnospiraceae* NK4A136 maintained high abundance independently on the administration of extracellular ASC oligomers **(Figure 7B)**. The decrease of pathobionts found in the in *Pycard*^-/-^ mice fed with a HFD was not affected by the administration of recombinant ASC oligomers, but interestingly the mucin-degrading bacteria *Akkermansia* tended to increase after extracellular ASC administration **(Figure 7B)**. We also observed a tendency to recover the abundance pattern of the different class of bacteria observed in *Pycard*^+/+^ mice, especially those less abundant classes, which were strongly reduced and responsible to decrease the biodiversity in *Pycard*^-/-^ mice after a HFD **(Suppl. Figure 6B)**. Additionally, ASC oligomer administration to in *Pycard*^-/-^ mice tended to recover the two most abundant families of bacteria (Akkermansiaceae and Lachnospiraceae) towards wild type mice abundance percentages **(Suppl. Figure 6C, D)**. This suggests that during HFD, inflammasome oligomers might be eliminated by the intestine where it could impact in gut microbiota composition, creating a beneficial environment in the gut by the lack of cellular free ASC oligomers. According to the increasing levels of mucin-degrading bacteria after treatment with extracellular ASC oligomers, we observed that this treatment also resulted in mucus not correctly attached between the *villi* space when compared with the *Pycard*^-/-^ mice without recombinant ASC oligomers administration **(Figure 7C)**. However, we did not observe differences in the transcript expression for the *ST6Galnac* gene family in the intestine **(Figure 7D)**.

**Figure 7.**
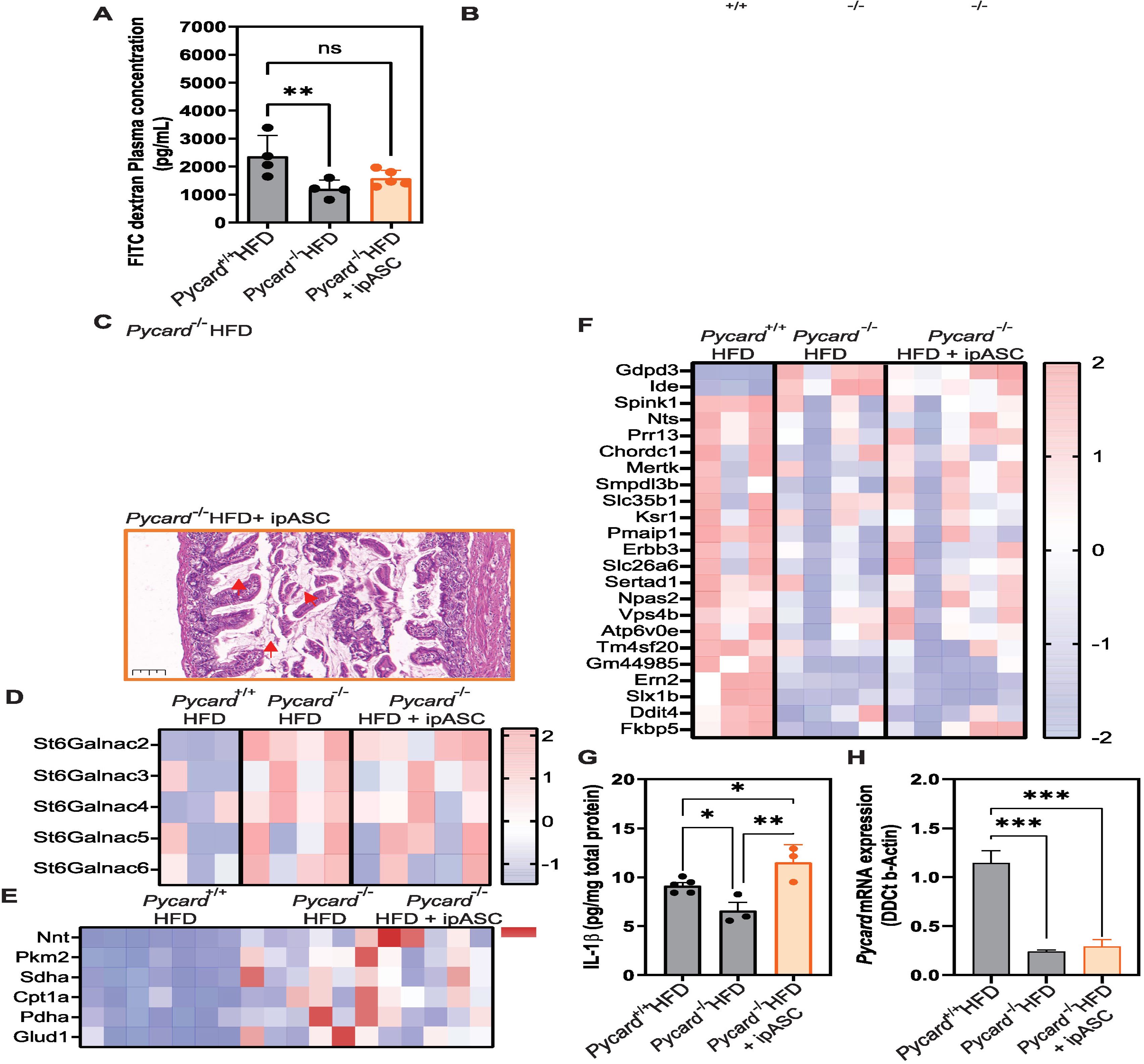
Extracellular ASC oligomers partially rescue intestinal microbiota and mucosal physiopathology during high fat diet. **A.** Model representation of the treatment with recombinant ASC oligomers in combination with high fat diet (HFD). **B.** Intestinal permeability measured by 4 kDa FITC-dextran in *Pycard*^-/-^ mice with intraperitoneal injection of recombinant ASC oligomers (+ipASC, orange, n=5) fed for 5 months with a HFD. Wild type (*Pycard*^+/+^, n=4) and *Pycard*^-/-^ (n=4) from Figure 3 are shown in grey for comparison. **C.** Relative abundance of *Odoribacter*, Lachnospiraceae NK4A136, Coriobacteriaceae UGC002, *Lactobacillus*, *Enteractinococcus, Atopostipes, Akkermansia*, *Streptococcus* and *Dorea* in the same conditions as in B. **D, E.** Haematoxylin and eosin histology of the small intestine of *Pycard*^-/-^ mice fed with a HFD for 5 months in the absence (D) or with ipASC (E). Scale bar: 200 μm; blue arrows indicate presence of mucus between *villi* and red arrows indicate not attached mucus. **F.** *ST6Galnac* gene family expression in the small intestine of mice treated as in B. **G.** Relative gene expression of *Nnt, Pkm2*, *Sdha*, *Cpt1a*, *Pdha*, and *Glud1* in the small intestine of mice treated as in B. **H.** Differential expressed genes in the small intestine of mice treated as in B. **I.** Interleukin (IL)-1β in the small intestine of mice treated as in B. **J.** Relative mRNA expression of *Pycard* in the small intestine of mice treated as in B. *Pycard*^+/+^ (n=3-5), *Pycard*^-/-^ (n=3-4), *Pycard*^-/-^ +ipASC (n=3-5). Each dot represents a distinct animal, bars represent the mean, error bars represent SEM. One-way ANOVA with multivariant Holm-Sidak’s test was used for B, C, G, I-J (ns, p>0.05; *p≤0.05; **p≤0.01; *** p≤0.001).

The administration of recombinant ASC specks in *Pycard*^-/-^ mice during HFD induced a partial reversion of the expression of some genes in the intestinal epithelia, especially those genes related to mitochondrial catabolism **(Figure 7E)**. Furthermore, bulk RNA-sequencing in the intestinal tissue revealed that extracellular ASC treatment of *Pycard*^-/-^ mice during HFD resulted in intermediate transcriptional profile between the wild type and the non-treated *Pycard*^-/-^ mice **(Figure 7F)**. As expected, intraperitoneal administration of recombinant ASC oligomers reverted IL-1β in the small intestine **(Figure 7G)**, confirming previous research where extracellular ASC oligomers could be internalized by macrophages and reconstitute inflammasome activity (8). We then confirmed that ASC gene expression was absent during the administration of recombinant ASC oligomers in the *Pycard*^-/-^ mice **(Figure 7H)**, confirming that the phenotype observed was independent of the endogenous expression of ASC.

ASC intraperitoneal treatment did not affect weight gain or glucose tolerance **(Suppl. Figure 7A, B)**. However, administration of ASC oligomers under HFD induced a tendency to increase the maximum peak of glycemia at 20 minutes after glucose injection in *Pycard*^-/-^ mice **(Suppl. Figure 7B)**. In addition, differentially to the effects observed in the small intestine, intraperitoneal administration of recombinant ASC oligomers did not rescue wild type phenotype of hepatic damage and fat accumulation after a HFD **(Suppl. Figure 7C, D)**, and similar low levels of hepatic IL-1β and high hepatic metabolism were also found after administration of ASC recombinant oligomers in the *Pycard*^-/-^ mice **(Suppl. Figure 7E, F)**. This suggests that extracellular ASC oligomers impact at the level of the small intestine, but not in hepatic physiopathology consequences, that might be controlled by the intracellular function of ASC-dependent inflammasomes or its downstream consequences.

Overall, ASC-dependent inflammasomes control metabolic syndrome induced by HFD at different levels, modulating intestinal permeability and inflammation, dysbiosis, glucose tolerance, hepatic inflammation, metabolism and steatosis. The presence of extracellular ASC oligomers partially modulates these effects but mainly impacted in dysbiosis and small intestine metabolism and inflammation.

## Discussion

In this study we found that ASC deficiency presented a protective effect during obesity induced metabolic syndrome. This protection was observed at different levels, including the decrease of intestinal permeability, dysbiosis, changes in the integrity, metagenomic, metabolism and physiology of the small intestine epithelia, and a decrease of IL-1β production. This protective effect was also found in hepatic metabolism, inflammation and steatosis. Dysbiosis, intestinal metabolism and IL-1β production were reverted in the *Pycard*^-/-^ mice when recombinant ASC oligomers were intraperitoneally administrated during the HFD, suggesting a differential role of extracellular ASC oligomers for microbiota and intestine and hepatic functions.

Our study is in line with a previous study where ASC was also found able to regulate glucose tolerance in obesity (27), but other studies found an ASC-independent protection of the intestinal mucosa and dysbiosis that were dependent on the mother and cage (28). In this study we found that obesity-induced dysbiosis was partially reverted by administration of extracellular ASC oligomers, and cohousing demonstrated that specifically the presence ASC influences dysbiosis during HFD. This suggests that certain differences in microbiota composition can be explained by the cage or the mother, but our results support changes in the intestinal microbiome dependent of the extracellular presence of ASC during HFD. Additionally, previous studies used a HFD composition of 60% Kcal fat content (27), and humans that ingest high fat in westernized diets does not usually reach 40% Kcal in fat consumption (29,30). Our model with a 45% Kcal in saturated fat content is focused to mimic the human physiopathology of obesity and could contribute to potential differences with other studies.

Intestinal permeability has been related to the composition of the gut microbiota (31), and our results supported this idea as HFD impact intestinal microbiota biodiversity. HFD increase the abundance of species of the Verrucomicrobiae class, including *Akkermansia* genus, and specifically *A. muciniphila,* a mucin-degrading bacteria which normally is beneficial for the gut, but in excess, it can dysregulate gut homeostasis (32–34). However, the pathological consequences of HFD could not be attributed exclusively to *Akkermansia* abundance. It is reported that gut pathobionts have an advantageous niche when highly increased *Akkermansia* degrades mucus (35). Our results revealed that endogenous ASC was associated with the growing of *Dorea,* which could be considered a pathobiont (36). Also, we observed *Streptococcus* increase in an ASC-dependent manner, but the relationship between *Streptococcus* genera and the host is more complex and depends on the specie studied (37–39). Moreover, the decrease of *Akkermansia* in the gut create a new niche that can be colonised other by less abundant bacteria, reducing dysbiosis and increasing the microbial biodiversity as we observe for the *Pycard*^-/-^ HFD fed mice group. The decrease in mucus production could reduce the attachment of segmented filamentous bacteria, which then induces an immune response to maintain homeostasis of the intestinal mucosa (40). In our study, we also found high abundance of *Odoribacter* in ASC deficient mice, which is known to control glucose levels and inflammation (41). The protection of dysbiosis by the deficiency of ASC was also corroborated at histological level, where lack of ASC prevented the inclusion of bacteria in the intestinal mucosa, with an increase of mucus between the intestinal *villi*. Additionally, administration of recombinant ASC oligomers rescued the phenotype of ASC-deficient mice at intestinal level and promoted a dysbiosis similar to the wild type mice during HFD, reducing also intestinal mucus layer. Indeed, intestinal gene expression showed an increase of the *ST6Galnac* gene family in ASC deficient mice, being these genes codifying for a family of sialyl-transferases which induces mucus sialylation and promoting intestinal mucosa protection from bacterial degradation (42). Especially, we observe a partial reversion of *ST6Galnac6* expression after administration of ASC oligomers, which is specifically important to protect the mucosa from bacterial invasion (43). Concomitantly with this effect, we also found that ASC controlled intestinal epithelia gene expression related with the innate immune response against pathogens, inflammation, and epithelial integrity, some of them are considered as markers for obesity and cancer (44,45). Moreover, the increasing expression of the insulin degrading enzyme (*Ide*) gene that we observed in *Pycard*^-/-^ mice under HFD could explain the increasing tolerance to glucose, as this enzyme protects from insulin resistance, and obesity in response to high glycemia levels (46,47). Additionally, we found that the intestinal production of IL-1β dependent on ASC might be attributed to the NLRP3 inflammasome activation, as previous studies have found the participation of NLRP3 during obesity (17,48). However, we did not discard the participation of other inflammasomes, which are also dependent on ASC and expressed in intestinal epithelia, that could contribute to IL-1β production (49–51).

ASC oligomers have been detected extracellularly in different pathologies, including autoinflammatory syndromes, amyloidosis, Alzheimer disease, and HIV infection (8–13,26). However, we found that obesity, glucose tolerance and liver steatosis, long term pathological consequences of metabolic syndrome (52,53), were not affected by the presence of extracellular ASC oligomers, which being probably dependent on the intracellular functions of ASC. In fact, we found an increase of peritoneal cells in the absence of ASC, probably as consequence of the reduced cell death of peritoneal cells upon ASC-dependent inflammasome activation (54). Therefore, the protective effect of ASC depletion could be mediated by a mechanism of regulation which depends on the inflammasome induced ASC oligomerization, being the release of these oligomers controlling additional pathological consequences during obesity. Recent developments of nanobodies-targeting extracellular ASC oligomers (12) might be beneficial to treat dysbiosis associated to HFD. Additionally, metabolic syndrome could be controlled by conventional antibodies targeting ASC or other small molecules blocking ASC oligomerization have been shown to reduce inflammation in different models (55–59). This is of relevance, as multiple inflammasomes have been involved in the pathophysiology of metabolic syndrome (49,60–62), and targeting ASC could modulate all of them with a single drug that could go beyond inflammasome or IL-1 blocking therapies (63,64). Particularly, we found that small intestinal cells presented an exacerbated catabolic pathway in *Pycard*^-/-^ mice. This might be due because of the increased glucose tolerance with an augmented glycolytic pathway and TCA through the enzyme’s pyruvate kinase and pyruvate dehydrogenase, while decreasing anaerobic lactate dehydrogenase activity. This regulation of glycolysis would reduce fat accumulation in the liver and might increase hepatic metabolism (65). The increased metabolism found in the liver could be the consequence to reduce fat accumulation, in a very similar manner as occurs for the small intestine. However, differentially to the small intestine, extracellular ASC oligomers did not rescue the pathological phenotype of the liver.

In summary, we report a mechanism of protection to the hyperglycaemic, westernized and high-saturated fat diets which is dependent on the deficiency of the inflammasome adaptor protein ASC. This mechanism is characterized by an exacerbated glucose metabolism together an increasing inflammatory phenotype and a decreasing intestinal permeability by the prevention of dysbiosis and mucus degradation. Extracellular ASC have a specific role on the development of the mucosal immunology and dysbiosis during HFD, but not in the systemic consequences of metabolic syndrome, including hepatic function, tolerance to glucose, and weight gain, which appear to be controlled by the intracellular ASC function forming inflammasomes. Further studies are warranted to evaluate potential therapeutic approximations based in the inhibition of ASC oligomerization.

## Materials and Methods

### Animal Models

Animal studies were approved by the Animal Health Service of the General Directorate of Farming, Fishing and Aquiculture of the Council of Murcia (CARM, ID# A13211201) once were positively evaluated by the Animal Experimentation Ethics Committee (CEEA) of the University of Murcia (#738/2021) according to the 53/2013 Royal Decree and the 32/2007 National Law. Mice were bred in specific pathogen-free conditions with a 12:12 h light-dark cycle in the animal facility from the University of Murcia, according to governmental guidelines 86/609/CEE. Age and gender-matched ASC-deficient (*Pycard*^-/-^) and wild type C57BL6/J (*Pycard*^+/+^) backgrounds mice were provided to a free access to a standard laboratory chow diet. For the obesity model, mice were fed with a 45% saturated fat HFD (D12491, Research Diets). Mice were autopsied after euthanasia with excess of isoflurane and cervical dislocation according to the ethical protocols of the national and international animal experimentation guidelines. After sacrifice, feces, peritoneal lavage, the right lobule of the liver, 1 cm of the proximal small intestine and blood samples were recovered. All samples were recovered strictly in the morning between Zeitgeber Time (ZT)0 and ZT4, to avoid the interference of the circadian changes. Histological samples were collected on paraformaldehyde 4% (Sigma-Aldrich, Burlington, MA) at room temperature. For RTqPCR samples, tissues were collected on RNA later (Qiagen, Germany) and frozen on liquid nitrogen. Feces were collected in a dry tube and frozen on liquid nitrogen. Peritoneal cells were collected on Phosphate Saline Buffer (PBS) and kept at 4°C before plating. Plasma was collected in heparin tubes (Microvette® CB 300 LH, Sarstedt, Germany). For tissue storage, samples were stored at -80°C.

### In vivo treatments

ASC Oligomers were produced and purified as we previously described in Alarcón-Vila et al. 2024 (11). Counted ASC oligomers were diluted into 1x PBS at a concentration of 4×10^6^ oligomers per 100μl, ready to use it *in vivo*. ASC Oligomers were weekly injected intraperitoneally at a concentration of 4×10^6^ oligomers per mouse. Glucose Tolerance Test (GTT) was done at 3 and 5 months after the administration of the HFD upon gender and sex matched animals. Prior the GTT assay, animals were fasted the day before for 14 hours to avoid food interference on the glucose levels. Basal fasted glycemia was measured before glucose administration. Following, 1g/Kg of D-glucose (Sigma-Aldrich) was intraperitoneally injected to each mouse, and glycemia was measured every 20 minutes for a minimum of one hour. The last point was measured at 90 minutes after glucose injection. At the end of the experiment, mice were fed back again. For the intestinal permeability assay, mice were deprived of drinking water during one hour at ZT0. Immediately, 4 kDa FITC-dextran were administered orally by gavage at a concentration of 600 mg/Kg of mouse. After one hour, blood was collected, and plasma was obtained by centrifugation at 2000 x g for 8 minutes. Fluorescence at 450 nm excitation and 570 nm emission was measured from the plasma using a Synergy Mx plate reader (BioTek).

### Cells and Treatments

5×10^5^ peritoneal cells (lymphocytes were not included into the count) were seeded per well in 24 wells plate at 1 million/mL concentration in RPMI medium (Sigma Aldrich) supplemented with 10% fetal bovine serum (Biowest), 2 mM GlutaMAX (Thermo Fisher Scientific) and 1% Penicillin Streptomycin (Corning). After overnight incubation at 37°C and 5% CO_2_ non-adherent cells were removed and LPS from *Escherichia coli* 055:B5 (Sigma-Aldrich) was added at 1 μg/mL for 4 hours in OptiMEM (Gibco). After priming, supernatant was removed and nigericin (Sigma-Aldrich) was added at 10 μM in a buffer containing 147 mM NaCl, 10 mM HEPES, 13 mM D-glucose, 2 mM CaCl_2_, 1 mM MgCl_2_, and 2 mM KCl (pH 7.4) for 30 minutes. MCC950 (Sigma-Aldrich) was added simultaneously to the nigericin at 10 μM.

### Tissue lysates preparation

Liver and small intestines previously stored at -80°C were mechanically lysed and kept on ice. After being centrifuged at 16,000 x g for 10 minutes, supernatant was recovered, and protein concentration was determined using the Bradford reagent (Sigma Aldrich).

### ELISA determinations

After *in vitro* peritoneal cell stimulation, supernatants from the cells were collected and centrifuged at 16,000 x g for 30 seconds. IL-1β and IL-6 were measured in the peritoneal cells supernatants, as well as in the liver and small intestine soluble fractions, using the mouse Quantikine^TM^ ELISA kits, catalogue number MLB00C-1 and M6000B respectively (R&D Systems, Minneapolis, MN) following the manufacturer indications, and results were analysed in a Sinergy MX plate reader (BioTek, Winooski, VT).

### Histological samples preparation and analysis

Fixed liver and small intestine were processed, paraffin-embedded and sections stained with haematoxylin and eosin using standard methods. Slides were examined using a Zeiss Axio Scope AX10 microscope with an AxioCamICC3 (Carl Zeiss, Germany).

### RNA extraction and RTqPCR determination

RNA was isolated using the RNeasy mini kit according to the manufacturer instructions (Qiagen). Cell lysis was performed with the addition of RLT buffer supplemented with β-mercaptoethanol. RNA integrity and concentration were assessed using the NanoDrop spectrophotometer (Thermo Fisher Scientific, Waltham, MA) Isolated RNA was reverse transcribed with the iScript™ cDNA Synthesis Kit (BioRad, Hercules, CA) according to the manufacturer’s instructions. The resulting complementary DNA (equivalent to 500 ng of total RNA) was amplified using the mix SYBR Premix ExTaq (Takara) and detected at real time on the iCyclerMyiQ thermocycler (BioRad). Quantitative PCR was performed using forward and reverse primers (**Supplemental Table 1**). On completion of the PCR amplification, a DNA melting curve analysis was carried out to confirm the presence of a single amplicon. *Actb* gene was used as a reference gene to normalize the transcript levels. Relative mRNA levels (2^-ΔΔCt^) were determined by comparing the ΔCt values for treated and control groups (ΔΔCt).

### RNA library preparation and sequencing

RNA libraries were prepared using the QuantSeq 3’mRNA-Seq V2 Library Prep Kit REV with UDI Set B (Lexogen, Austria), following the manufacturer’s protocol. Briefly, 500 ng of total RNA per sample was used as input for poly(A) mRNA capture using oligo-dT beads. Captured mRNA was fragmented and reverse-transcribed into cDNA. Second-strand synthesis and adapter ligation were performed to generate double-stranded cDNA libraries. Libraries were indexed with 12-nucleotide unique dual indices for multiplexing and amplified using polymerase chain reaction (PCR). The quality and quantity of the final libraries were evaluated using the Agilent 2100 Bioanalyzer and the Qubit 4 Fluorometer (Thermo Fisher Scientific). Libraries were sequenced on a NextSeq 2000 system (Illumina, San Diego, CA) in paired-end mode with a read length of 2 × 100 bp to a target depth of 30 million reads per sample.

### RNA sequencing analysis

For RNAseq-data processing and alignment, raw sequencing data were processed using the bcl2fastq software (Illumina) to demultiplex and convert base call files into FASTQ format. Quality control of the FASTQ files was performed using FastQC (v0.11.9) to assess read quality, adapter content, and GC distribution. Low-quality reads and adapter sequences were trimmed using Trimmomatic (v0.39). Cleaned reads were aligned to the mouse reference genome (GRCm39) using STAR (v2.7.10a) with default parameters, generating BAM files for each sample. For the differential gene expression, Gene expression quantification was performed using featureCounts (v2.0.3) to obtain raw read counts per gene, based on the GENCODE M28 annotation. Count data were provided in a tab-delimited file containing unique read counts for each gene across samples. The dataset was preprocessed using R (R Core Team, 2023). Gene count matrices were normalized using the DESeq2 package (66), which applies a variance-stabilizing transformation to adjust for sequencing depth and RNA composition biases. Differential gene expression analyses were performed using DESeq2’s generalized linear model framework, considering the experimental condition as the primary variable. To identify the top differentially expressed genes, the normalized counts were z-score transformed. Heatmaps of the top 20 differentially expressed genes were generated using the pheatmap package (67), with sample group annotations included for clarity. Gene identifiers were converted to gene symbols using the org.Mm.eg.db annotation database (68). Pathway enrichment analysis was performed on differentially expressed genes using the Cluster Profiler package (v4.8.1) in R. Gene Ontology (GO) biological processes were analyzed, with pathways considered significant at a false discovery rate < 0.05. The selected resulting pathways were visualized using dot plots generated with ggplot2 (v3.4.0).

### Bacterial DNA extraction

Fecal samples were collected directly from the bedding-free cage floor to avoid contamination. Immediately after collection, fecal samples were frozen at −80 °C until further processing. Bacterial DNA from feces was extracted using the Purelink^TM^ Microbiome DNA Purification Kit (Thermo Fisher Scientific), according to the manufacturer instructions. The concentration and purity of extracted DNA were assessed with a NanoDrop spectrophotometer (Thermo Fisher Scientific).

### 16S ribosome RNA gene amplification, and sequencing

For the amplification of the 16S rRNA gene, primers targeting the V3-V4 hypervariable regions were designed to cover the bacterial domain (84.2%), archaea (75.8%), based on the SILVA database (March 2020). PCR reactions were performed using 10 ng of DNA as input, combining targeted amplification with barcode addition for sample multiplexing. The resulting amplicons were approximately 606 base pairs in length. The amplification process utilized the Quick-16S Plus NGS Library Prep Kit (Zymo Research) according to the manufacturer’s instructions. Following amplification, libraries were pooled in equimolar volumes and purified using magnetic beads to remove residual primers and small fragments, as specified by the kit’s protocol. Sequencing of the libraries was performed on the NextSeq 2000 platform (Illumina, San Diego, CA) using a paired-end (2x300 bp) approach. This setup ensured high-resolution reads suitable for downstream taxonomic and functional analyses. A minimum of 50,000 reads per sample was targeted to achieve sufficient depth for robust microbial community profiling.

### Metagenomic analysis

The raw data was first quality controlled by trimming low-quality bases and removing adapters using Trimmomatic. Next, the sequences were denoised and chimeras were removed using the DADA2 algorithm implemented in QIIME 2. Following denoising, the sequences were assigned taxonomy at the genus level using the SILVA v138 database and a Naïve Bayes classifier, which classifies each ASV based on its sequence similarity to known reference sequences. Phyloseq package was used to create a phyloseq object from the ASV table, taxonomic classification, and metadata. The data was then aggregated at the genus level using the tax_glom() function, and relative abundances were calculated using the transform_sample_counts() function in the phyloseq package. The entire workflow was performed using the phyloseq, ggplot2, dplyr, and writexl R packages (69,70).

### Statistical analysis

For two variables analysis, normality tests Shapiro-Wilk and Kolmogorov-Smirnov were used to elucidate parametric and non-parametric groups. Where at least, one of the independent groups was not following normality, non-parametric analysis was used. For testing differences between two independent groups, Mann-Whitney test was used for non-parametric data, and student *t*-test was used for parametric data. one-way ANOVA test was used for parametric data. For three or more variables analysis, two-way ANOVA test was used. Between each ANOVA test, multivariant test analysis was used to compare each bidirectional comparison between single groups (Holm-Sidak’s test for the parametric one-way ANOVA multivariant analysis, Sidak’s test for two-way ANOVA multivariant analysis). For correlations, non-parametric Spearman test was used. Outliers were discarded when were significant for the Grubb’s test.

## Abbreviations list

ASC: Apoptosis-associated Speck-like protein containing a CARD
cDNA: Complementary Deoxyribonucleic Acid
DEG: Differential Expressed Gene
DNA: Deoxyribonucleic Acid
FDR: False Discovery Rate
GO: Gene Ontology
GSDMD: Gasdermin D
GTT: Glucose Tolerance Test
HFD: High-Fat-Diet
IL: Interleukin
LPS: Lipopolysaccharide
mRNA: Messenger Ribonucleic Acid
NCD: Normal-Chow-Diet
NIN1: Ninjurin-1
NLR: Nucleotide-binding domain Leucine-rich Repeat receptor
NLRP: Nucleotide-binding domain Leucine-rich Repeat Pyrin domain-containing receptor
NOD: Nucleotide-binding and Oligomerization Domain
PCA: Principal Components Analysis
PCR: Polymerase Chain Reaction
PBS: Phosphate Saline Buffer
RNA: Ribonucleic Acid
TCA: Tricarboxylic Acids cycle
TLR: Toll-Like Receptor
ZT: Zeitgeber Time.

## Acknowledgements

We thank AI Gomez for cell culture and lab assistance, and the members of Dr. Pelegrin’s laboratory for all comments and suggestions. We also acknowledge to the animal facility and cell culture facility from the University of Murcia, as well as the SPF-animal house, Pathology, Genomics and Bioinformatics platforms from the Pascual Parrilla-Biomedical Research Institute of Murcia (IMIB).

## Authors Contribution

JJM-G visualized and administered the research. JJM-G and PP conceptualized, supervised, provided funding, and wrote the manuscript. JJM-G and SR did the formal analysis, wrote, and edited the manuscript. AB-M provided funding and supervision. JJM-G, AG, IQ-R, SV-M. CM-L and LH-N performed the experimental research.

## Supplemental information

### Resource table

Resource table.
| Reagent | Source | Identifier |
| --- | --- | --- |
| **Samples** |  |  |
| C57BL/6J (*Pycard*<sup>+/+</sup>) mice (M. musculus) | Jackson Laboratories | RRID: IMSR_JAX:000664 |
| ASC deficient mice in C57BL/6J background (*Pycard*<sup>-/-</sup>) | Mariathasan et al. 2004 (71) | N/A |
| ASC Oligomers | Alarcón-Vila et al. 2025 (11) | N/A |
| **Commercial** |  |  |
| D-Glucose | Sigma Aldrich | Cat. G7021 |
| 4Kda-FITC Dextran | Sigma Aldrich | Cat. 68059 |
| *Escherichia coli* LPS 055:B5 | Sigma Aldrich | Cat. L2880 |
| Nigericin | Sigma Aldrich | Cat. N7143 |
| MCC950 | Sigma Aldrich | Cat. 5.38120 |
| 45% High-Fat-Diet | Research Diets | Cat. D12491 |
| 32% Paraformaldehyde | Sigma Aldrich | Cat. P6148 |
| RNA later | Qiagen | Cat. 76106 |
| Bradford Reagent | Sigma Aldrich | Cat. B6916 |
| RTqPCR Oligomers | Sigma Aldrich | Suppl. Table 1 |
| Mouse IL-6 ELISA Kit - Quantikine | R&D Systems | Cat. M6000B |
| Mouse IL-1b ELISA Kit - Quantikine | R&D Systems | Cat. MLB00C1 |
| RNeasy mini kit | Qiagen | Cat. 74104 |
| iScript™ cDNA Synthesis Kit | BioRad | Cat. 1708890 |
| SYBR Premix ExTaq | Takara | Cat. RR420A |
| QuantSeq 3' mRNA-Seq V2 Library Prep Kit REV with UDI Set B | Lexogen | Cat. 225 |
| Purelink™ Microbiome DNA Purification Kit | Thermo Fisher Scientific | Cat. A29790 |
| Quick-16S Plus NGS Library Prep Kit | Zymo Research | Cat. D6432 |
| E-Total | Martinez-Garcia et al. 2019 (72) | N/A |
| RPMI-1640 Medium liquid without L-glutamine | Sigma Aldrich | Cat. R0883 |
| OptiMEM reduced Serum Media | Gibco | Cat. 51985-026 |
| GlutaMAX | Thermo Fisher Scientific | Cat. 35050038 |
| Fetal Bovine Serum (FBS) | Biowest | Cat. S181A |
| Penicillin-Streptomycin | Corning | Cat. 30-002-CI |
| **Software and packages** |  |  |
| GraphPad Prism version 9 | GraphPad Software Inc | https://www.graphpad.com/ |
| bcl2fastq software | Illumina | https://support.illumina.com/sequencing/sequencing_software/bcl2fastq-conversion-software.html |
| FastQC (v0.11.9) | Brabraham Bioinformatics | https://www.bioinformatics.babraham.ac.uk/projects/fastqc/ |
| Trimmomatic (v0.39) | Usadellab | http://www.usadellab.org/cms/?page=trimmomatic |
| STAR (v2.7.10a) | Dobin et al. 2012 (73) | https://code.google.com/archive/p/rna-star/ |
| featureCounts (v2.0.3) | Subread | https://subread.sourceforge.net/featureCounts.html |
| R Software | R Core Team | https://www.r-project.org/ |
| ClusterProfiler package (v4.8.1) | Bioconductor | https://bioconductor.org/packages/devel/bioc/html/clusterProfiler.html |
| ggplot2 (v3.4.0) | Tidyverse | https://www.tidyverse.org/blog/2022/11/ggplot2-3-4-0/ |
| pheatmap package | Kolde et al. 2019 (67) | https://cran.r-project.org/web/packages/pheatmap/index.html |
| phyloseq package | McMurdie et al. 2013 (70) | https://joey711.github.io/phyloseq/install.html |
| **Devices** |  |  |
| CentriVet GK Glucometer | ACON Laboratories Inc. | N/A |
| Synergy Mx Plate Reader | Biotek | N/A |
| Zeiss Axio Scope AX10 | Carl Zeiss | N/A |
| AxioCamICC3 | Carl Zeiss | N/A |
| NanoDrop | Thermo Fisher Scientific | N/A |
| iCyclerMyiQ thermocycler | BioRad | N/A |
| Agilent 2100 Bioanalyzer | Agilent | N/A |
| Qubit 4 Fluorometer | Thermo Fisher Scientific | N/A |
| NextSeq 2000 | Illumina | N/A |

**Supplemental Table 1.**
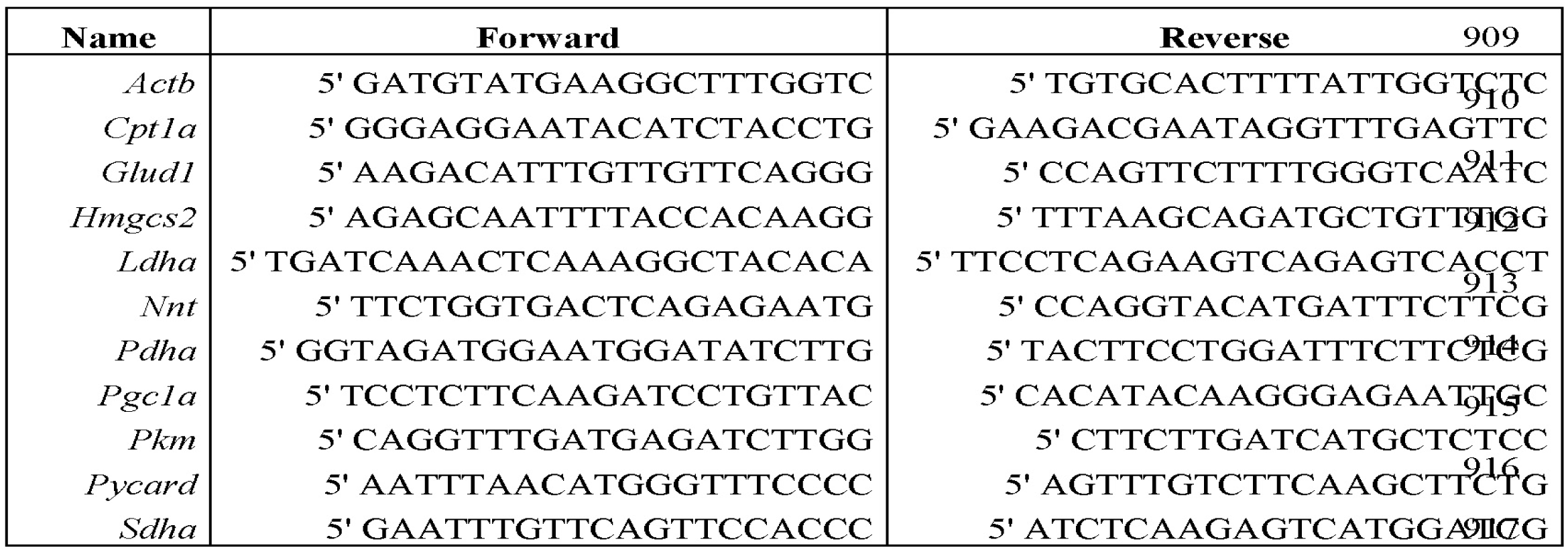
RT-qPCR primers sequences used in this study.

## Supplemental Figure Legends

**Supplemental Figure 1.**
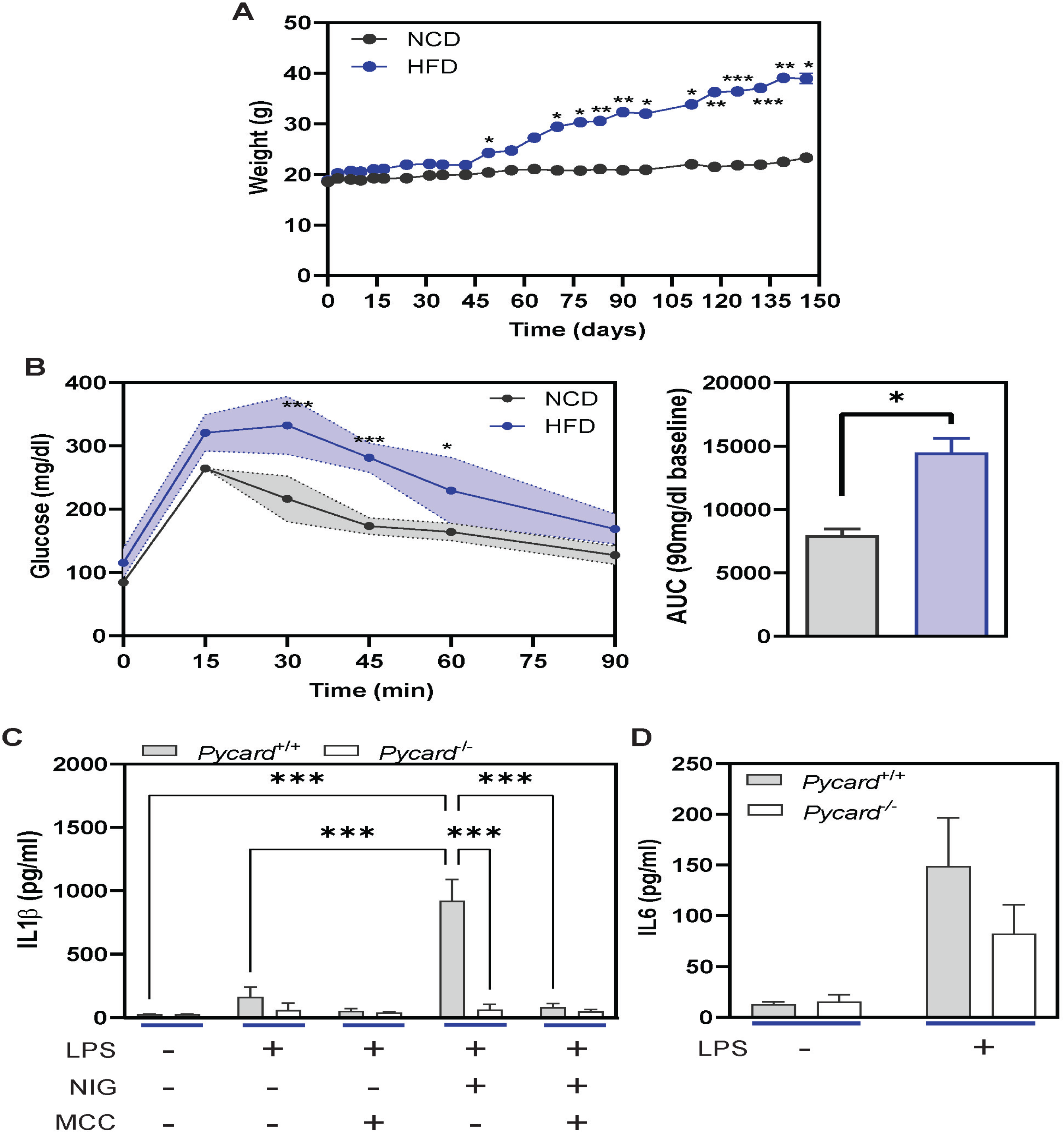
High fat diet increases glucose tolerance and body weight. **A.** Body weight on wild type (*Pycard*^+/+^) mice fed for 5 months under normal chow diet (NCD, grey, n=4) or high fat diet (HFD, blue, n=5). **B.** Glucose tolerance (left) and area under the curve (AUC, right) in wild type mice treated as in A. **C, D.** Interleukin (IL)-1β (C), and IL-6 (D) release from peritoneal cells collected from wild type (*Pycard*^+/+^, n=4, grey) or *Pycard*^-/-^ (n=4, white) mice fed for 5 months with a NCD and in vitro stimulation as indicated in the figure (LPS: lipopolysaccharide; NIG: nigericin, MCC: MCC950). Columns represent the mean of each condition and error bars represent SEM. Two-way ANOVA with multivariant Sidak’s test was used for A, B (left panel) and C. Mann-Whitney test was used for B right panel (ns, p>0.05; *p≤0.05; **p≤0.01; *** p≤0.001).

**Supplemental Figure 2.**
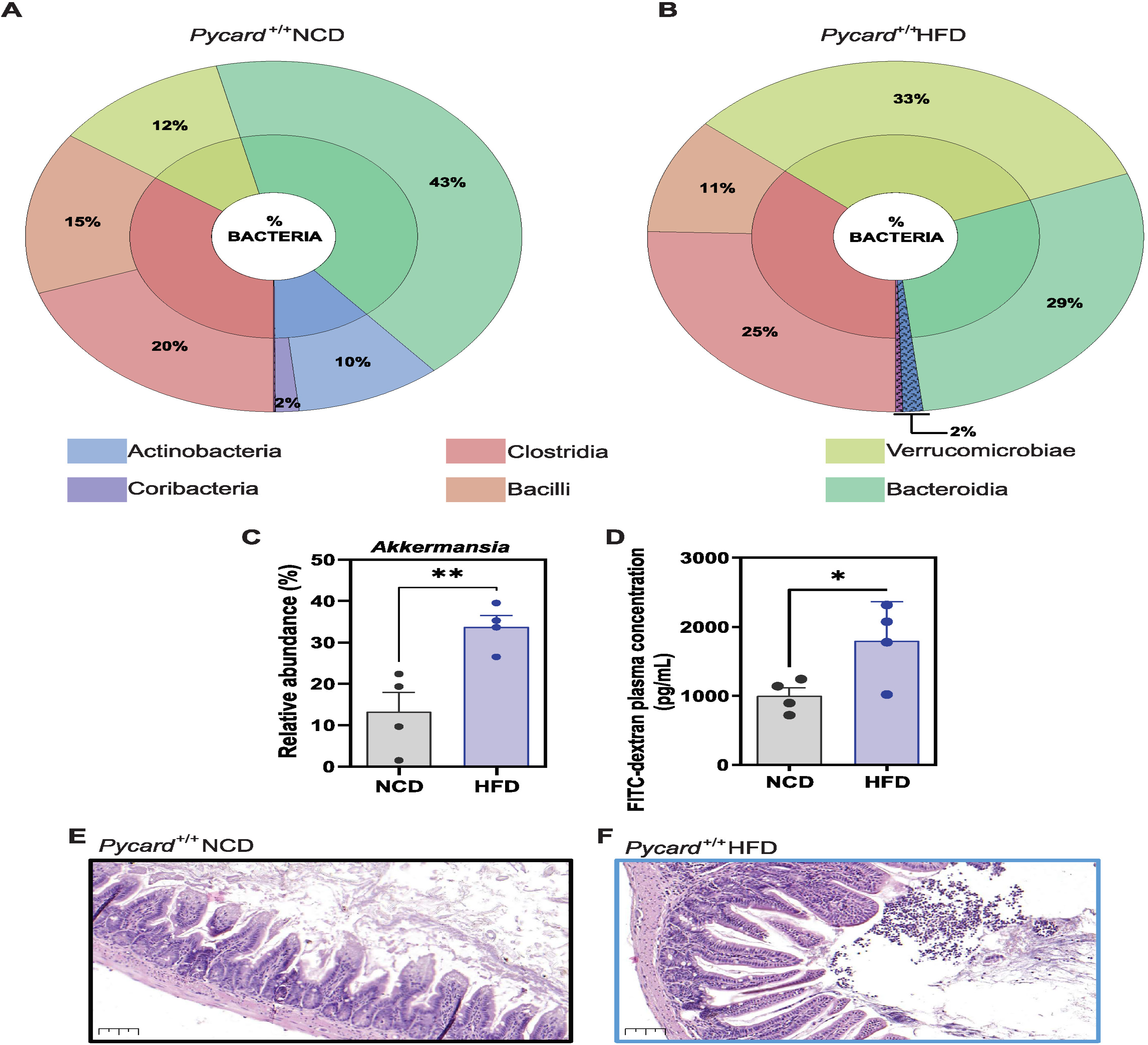
HFD induces dysbiosis and compromises gut epithelial protection. **A-C.** Relative abundance of the total faecal microbiota at class level (A, B), and of *Akkermansia* (C) on wild type (*Pycard*^+/+^) mice fed with a normal chow diet (NCD, n=4) in comparison with high fat diet (HFD, n= 5). **D.** Intestinal permeability measured by 4 kDa FITC-dextran in *Pycard*^+/+^mice treated as in A-C. **E,F.** Haematoxylin and eosin histology of the small intestine of *Pycard*^+/+^ mice fed for 5 months with a NCD representative of n=4 (E) or HFD representative of n= 5 (F). Scale bar: 200 μm. Each dot represents a distinct animal, bars represent the mean, error bars represent SEM. Unpaired *t*-test was used for C, D (*p≤0.05; **p≤0.01).

**Supplemental Figure 3.**
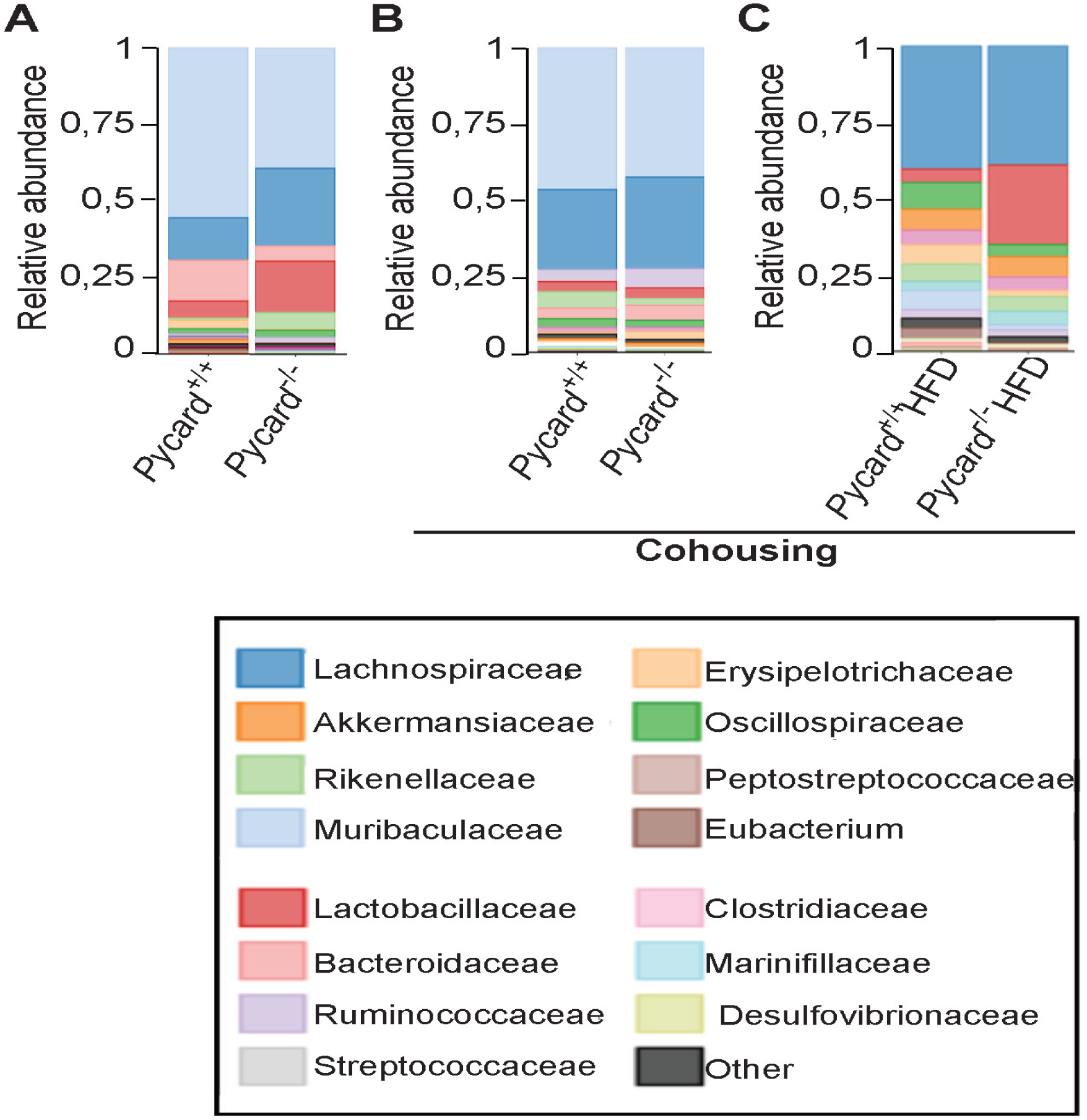
Cohousing partially restored wild type-dependent microbiota profile after HFD in *Pycard*^-/-^ mice. **A-C.** Relative microbiota abundance at family’s level of class on wild type (*Pycard*^+/+^, n=4) or *Pycard*^-/-^ mice (n=4) in normal chow diet (NCD) and genotype single housed (A) or cohoused (B) mice, and on cohoused mice after 3 months of high fat diet (HFD) (C).

**Supplemental Figure 4.**
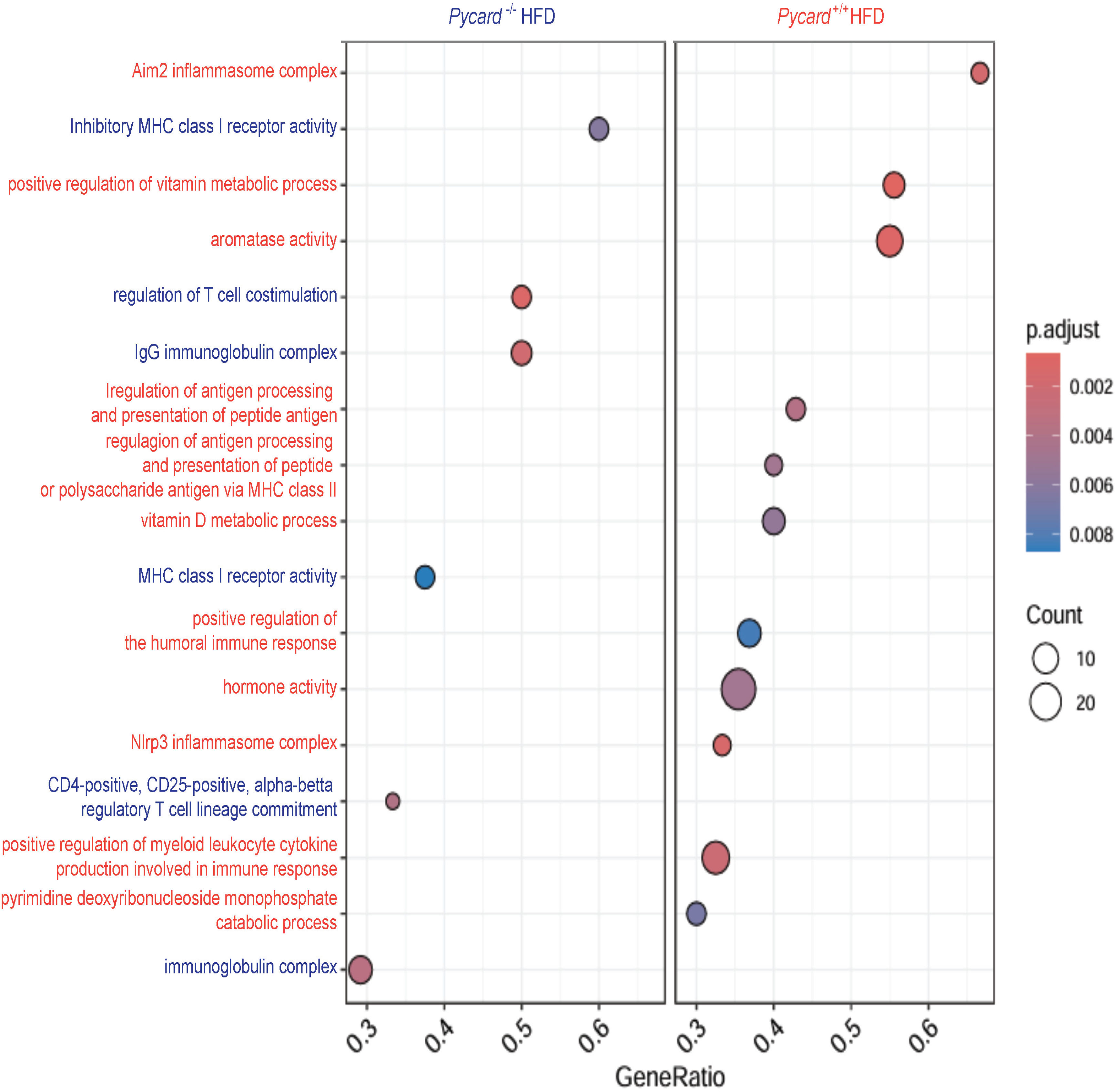
ASC deficiency increases regulation of inflammatory pathways in the small intestine during high fat diet. Extended vision of the Gene Ontology differential expressed pathways in the small intestine of wild type (*Pycard*^+/+^, right column) and *Pycard*^-/-^ (left column) mice fed for 5 months with a high fat diet (HFD). Color represents adjusted p value, and size the number of genes (count).

**Supplemental Figure 5.**
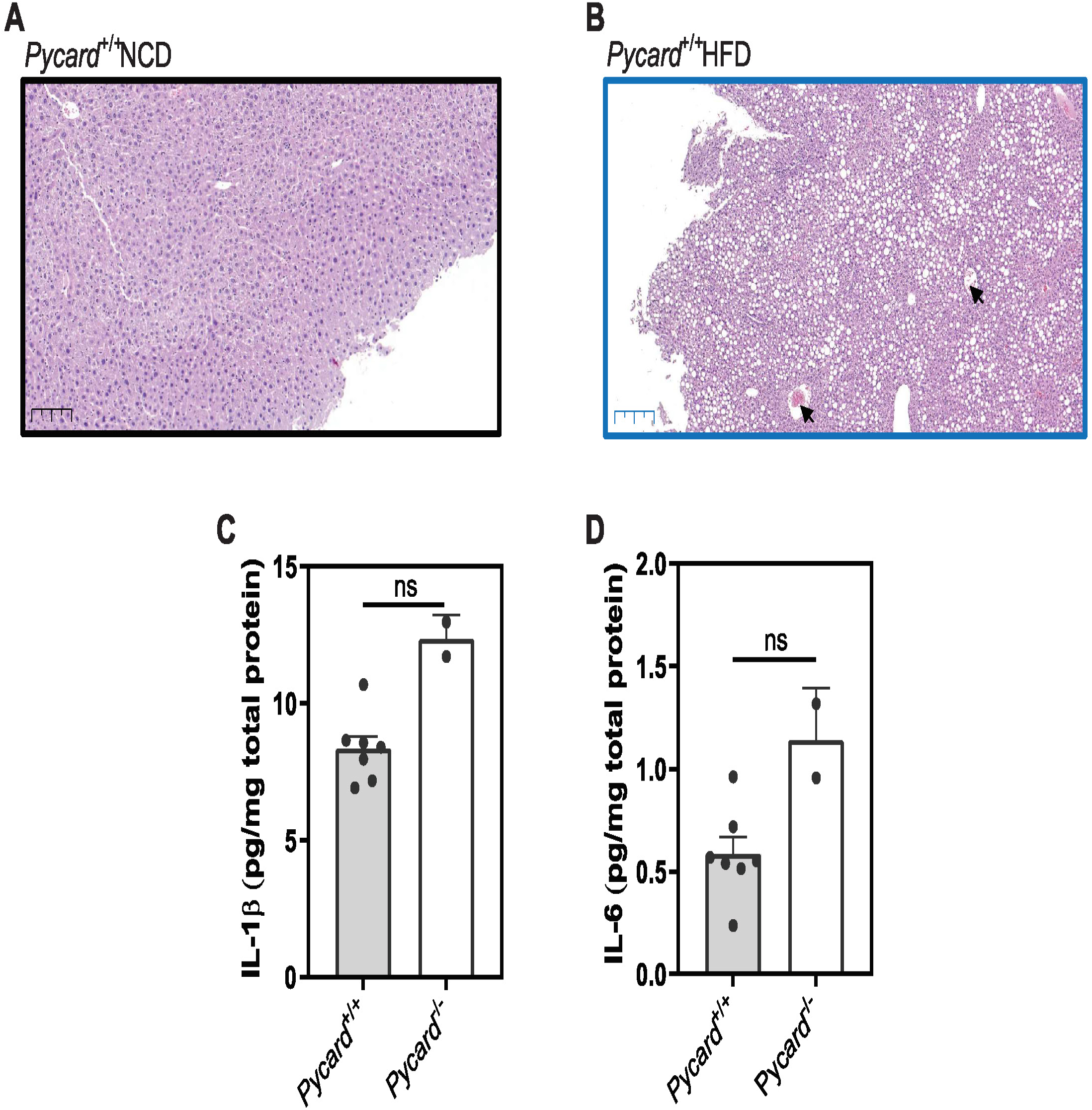
High fat diet induces liver steatosis and inflammation. **A, B.** Haematoxylin and eosin histology of the liver of wild type (*Pycard*^+/+^) mice fed for 5 months with a normal chow diet (NCD) (A) or a high fat diet (HFD) (B). Scale bar: 200 μm. **C, D.** Interleukin (IL)-1β (C) and IL-6 (D) in the liver of wild type (*Pycard*^+/+^, grey, n=7) or *Pycard*^-/-^ (white, n=2) mice fed for 5 months with an NCD. Each dot represents a distinct animal, bars represent the mean, error bars represent SEM. Mann-Whitney’s test (ns, p>0.05).

**Supplemental Figure 6.**
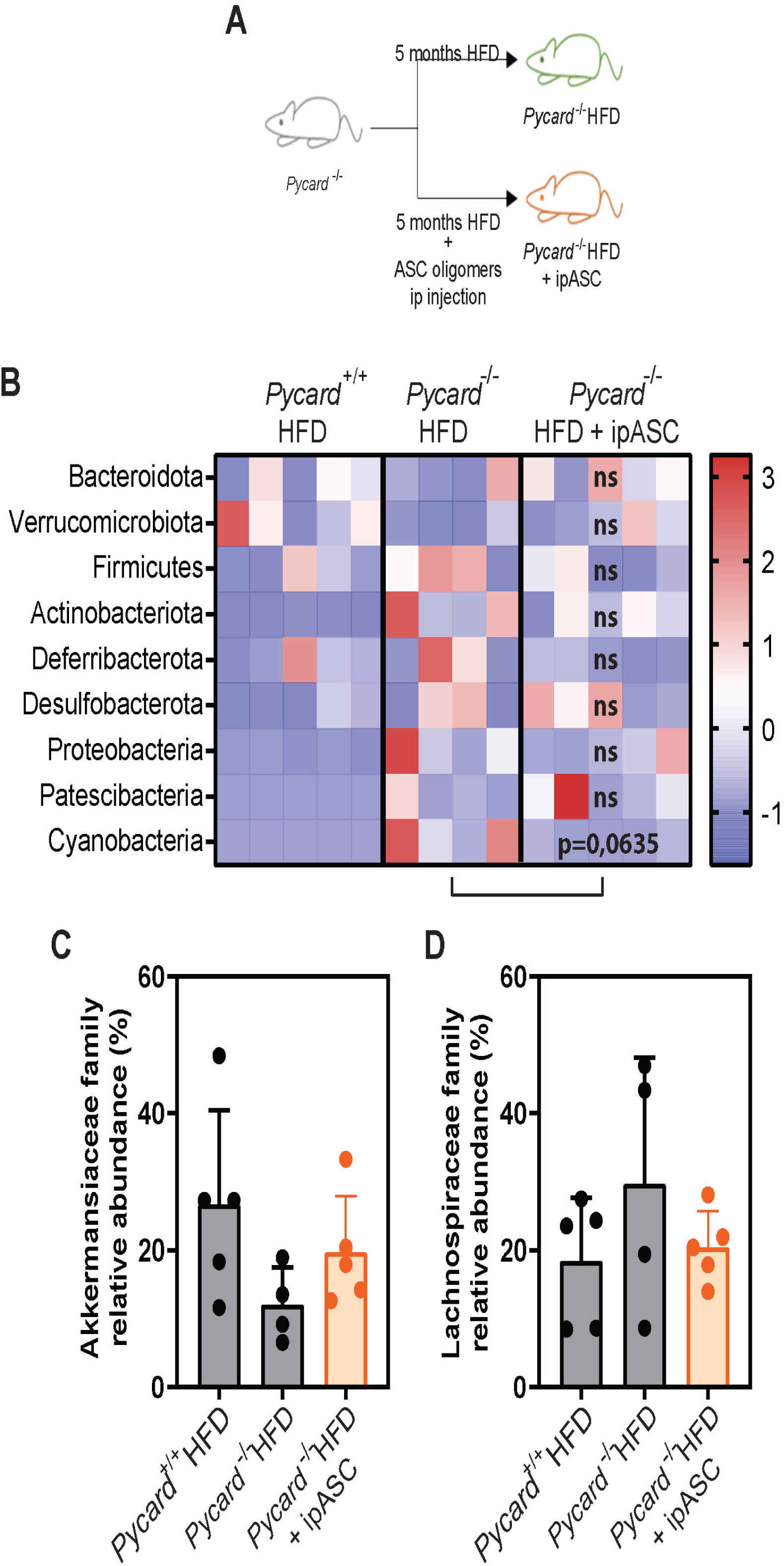
Intestinal microbiota after intraperitoneal administration of recombinant ASC oligomers during high fat diet. **A-C.** Relative microbiota abundance at class level (A), Akkermansiaceae (B) and Lachnospiraceae (C) families on feces of wild type (*Pycard*^+/+^, n=5), *Pycard*^-/-^ (n=4) or *Pycard*^-/-^ mice treated with intraperitoneal injection of recombinant ASC oligomers (ipASC, orange, n=4) fed for 5 months with high fat diet (HFD). Each dot represents a distinct animal, bars represent the mean, error bars represent SEM. One-way ANOVA with multivariant Holm-Sidak’s test, all comparison presented a p>0.05 (ns).

**Supplemental Figure 7.**
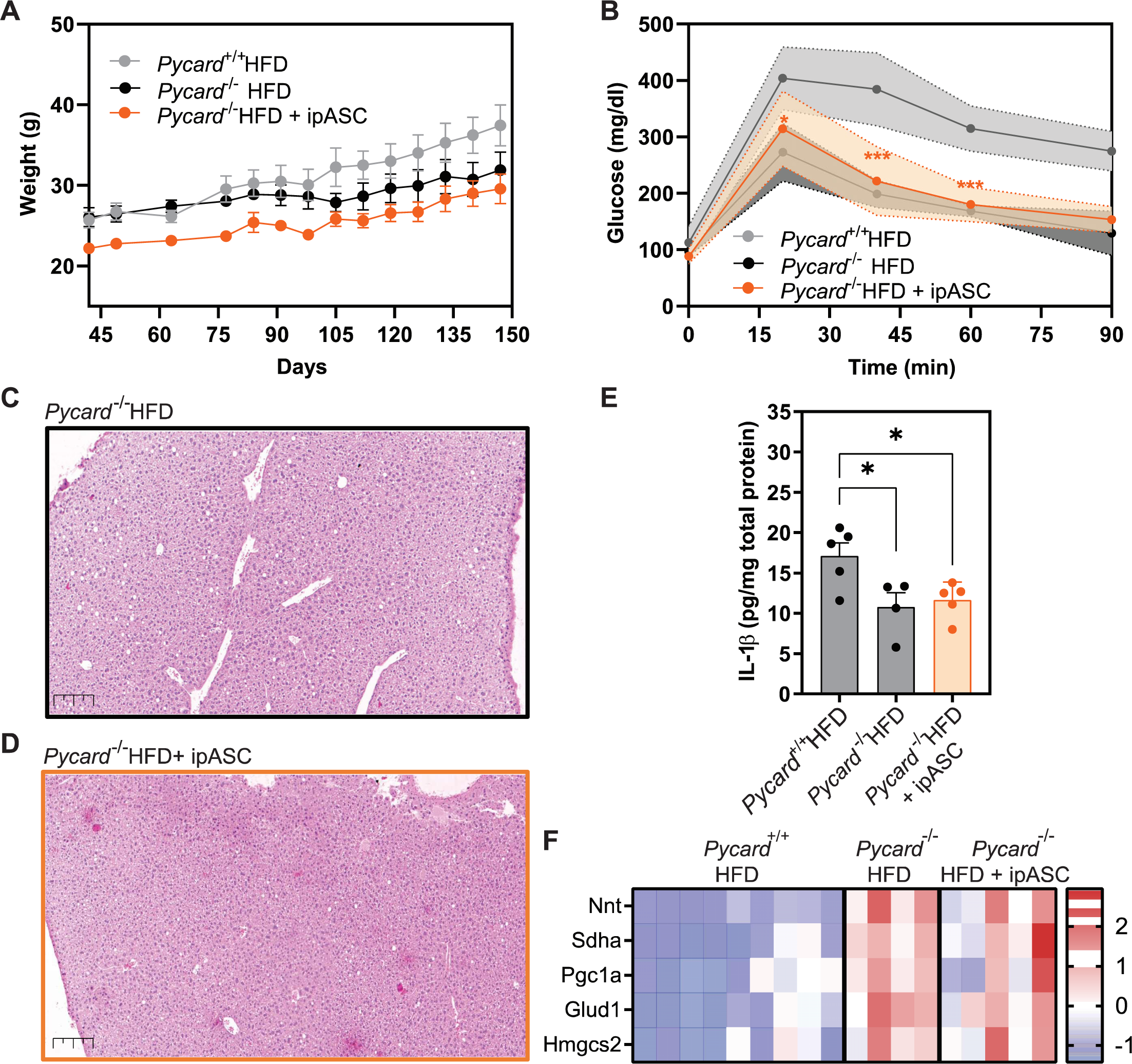
Intraperitoneal administration of recombinant ASC oligomers did not affect glucose tolerance, peritoneal inflammation or hepatic damage. **A.** Body weight from wild type (*Pycard*^+/+^, grey, n=5), *Pycard*^-/-^ (black, n=4), and *Pycard*^-/-^ with intraperitoneal injection of recombinant ASC oligomers (+ipASC, orange, n=5) mice fed for 5 months with a high fat diet (HFD). **B.** Glucose tolerance in mice treated as in A. **C, D.** Haematoxylin and eosin histology of the liver of mice treated as in A. Scale bar: 200 μm. **E.** IL-1β in the liver from mice treated as in A. **F.** Relative hepatic gene expression in mice treated as in A. Each dot represents a distinct animal, bars represent the mean, error bars represent SEM. Two-way ANOVA and multivariant Sidak’s test was used for A-B. One-way ANOVA and multivariant Holm-Sidak’s test was done for G, H (ns, p>0,05; *p≤0.05; **p≤0.01; *** p≤0.001).

